# PPARγ, a key modulator of metabolic reprogramming, stemness and chemoresistance associated with retrodifferentiation in human hepatocellular carcinomas

**DOI:** 10.1101/2024.09.02.610533

**Authors:** Yoann Daniel, Claudine Rauch, Lucille Moutaux, Karim Fekir, Lise Desquilles, Luis Cano, Daniel Catheline, Servane Pierre, Agnès Burel, Camille Savary, Catherine Ribault, Claude Bendavid, Bruno Clément, Caroline Aninat, Vincent Rioux, Orlando Musso, Bernard Fromenty, Florian Cabillic, Anne Corlu

## Abstract

Human hepatocellular carcinomas (HCCs) with cancer stem cell (CSC) features are a subclass of therapeutically challenging cancers. We recently showed that retrodifferentiation of hepatic cancer cells into CSC-like cells leads to metabolic reprogramming and chemoresistance. The molecular mechanisms whereby differentiated cancer cells switch towards a CSC phenotype are poorly understood. By studying metabolic reprogramming associated with HCC cell plasticity, we identified an unsuspected role of peroxisome proliferator-activated receptor (PPAR)γ in hepatic CSC phenotype acquisition.

Gene expression and metabolic analyses performed throughout cell differentiation/retrodifferentiation process of human HepaRG and HBG-BC2 HCC cells show that metabolic reprogramming in hepatic CSCs is associated with fragmented mitochondrial network, decreased respiration, *de novo* lipogenesis, fatty acid oxidation, but increased glycolysis and lipid storage. Mitochondrial genes downregulated in HepaRG-CSCs are also downregulated in the STEM HCC subclass. While PPARα is the main isoform in differentiated hepatic cells, we find high PPARγ expression in hepatic CSCs. Accordingly, nuclear localization of PPARγ is detected in human HCC tumors and PPARγ^high^/PPARα^low^ expression is associated with the STEM HCC subclass and a poor outcome in human HCC cohorts. PPARγ silencing or/and inhibition of its target gene pyruvate dehydrogenase kinase 4 reactivates cell respiration, increases reactive oxygen species production and sensitizes hepatic CSCs to chemotherapy. Conversely, PPARα activation synergizes with chemotherapy to induce cell death.

Targeting PPARγ, a key regulator of metabolic reprogramming and stemness in hepatic CSCs, or modulating the PPARγ/PPARα balance that finely tunes the differentiation/retrodifferentiation process in HCC deserves further investigation for antitumor therapy.

**Implications heading and statement:** PPARγ, a key regulator of metabolic reprogramming and stemness in hepatic CSC, reduces oxidative phosphorylation and reactive oxygen species production, therefore contributing to HCC chemoresistance.

## Introduction

Large-scale genome sequencing studies unraveled the molecular heterogeneity of hepatocellular carcinomas (HCCs) leading to various molecular classifications(1–4). HCCs with stem cell features constitute a subclass of therapeutically challenging cancers(5). Hepatic cancer stem cells (CSCs) form an heterogeneous cell compartment as they could result from transformation of stem/progenitor cells, but also and probably more often, from the acquisition of stem properties by liver cancer cells(5,6). Indeed, any hepatic cell lineage can retrodifferentiate into CSC as a result of oncogenic transformation(5) and/or inflammatory signals(7). Such plasticity potential contributes to both intra- and inter-tumor heterogeneity of HCCs(5,8) and makes it difficult to eradicate the whole tumor with single-agent therapies that do not address the diversity of cancer cell populations, including quiescent or slow-proliferating CSCs(9).

Recently, it has become clear that cell retrodifferentiation and subsequent phenotypic conversion were linked to metabolic shifts(10,11). Consistent with these observations, recent classifications have integrated cell metabolism as a discriminant parameter among HCCs(12,13). HCCs with liver periportal-type (HNF4α-driven) or perivenous-type (β-catenin-driven and fatty acid addicted) metabolic features belong to the non-proliferative, well-differentiated HCC class(13,14). Interestingly, the Yang’s subclasses with low and intermediate metabolic activities (gluconeogenesis, amino acid, lipid and drug metabolism) have the worst prognosis and match the poorly differentiated Hoshida’s S1 and Désert’s ECM and STEM subclasses(3,12,13). Therefore, identifying and targeting cancer cell-specific metabolism is a promising strategy to improve HCC therapy. Several works have focused on metabolic characteristics of tumor cells and CSCs, but few have addressed metabolic reprogramming associated with cancer cell retrodifferentiation.

We previously modeled HCC retrodifferentiation steps in two human HCC cell lines that, under defined culture conditions, display the CSC-, bipotent progenitor- and mature hepatocyte-like phenotypes(7,11). We reported that retrodifferentiation of human tumor-derived hepatocytes to CSCs leads to an upregulated expression of pyruvate dehydrogenase kinase 4 (PDK4), which prevents pyruvate from feeding the tricarboxylic acid (TCA) cycle. This upregulation is associated with stemness features in human HCCs(11). Moreover, the PDK4 inhibitor dichloroacetate (DCA) improves the efficacy of chemotherapies against hepatic CSCs(11). Unexpectedly, our transcriptomic analyses revealed that hepatic CSCs derived from differentiated tumor-derived hepatocytes have an increased expression of peroxisome proliferator-activated receptor (PPAR)γ, a key metabolic transcriptional modulator.

PPARγ is a master regulator of adipocyte differentiation through the control of lipid uptake, synthesis and storage, as well as glucose uptake. It also regulates a broad range of cellular functions e.g., inflammatory response, metabolism and apoptosis in various cell types(15). Interestingly, it is a positive transcriptional regulator of PDK4 in adipocytes(16) and lung cancer cells(17). In the liver, the predominant isoform is PPARα, which plays a crucial role in the maintenance of energy balance(18,19). During stress or starvation, PPARα activation spares pyruvate for gluconeogenesis *via* increased levels of PDK4, while it stimulates fatty acid oxidation (FAO) to fulfill energy needs(19,20). Although uncommon in the liver, PPARγ mediates anti-inflammatory and anti-fibrotic functions and maintains lipid/glucose homeostasis and insulin sensitivity in pathological conditions(21). Regarding HCC, PPARα is proposed as a positive prognosis marker(22). By contrast the role of PPARγ is controversial. Indeed, although PPARγ was mainly found to inhibit cell proliferation and HCC metastases *in vitro* and in mice(23–26), recent studies reported pro-tumorigenic effects in HCCs(27–29). Likewise, in other cancers, many studies suggest antitumor effects of PPARγ, but protumor effects are also reported(30). Focusing on CSCs, PPARγ activation promotes the eradication of CSCs in leukemia, prostate and colorectal cancers by modulating CSC self-renewal and differentiation(31–33). Conversely, it maintains ERBB2-positive breast CSCs(34) and PPARγ agonists increase the incidence of colorectal, renal and bladder cancers(35,36). It is therefore crucial to clarify the role of PPARs according to cell types and/or cell differentiation stages in HCCs before considering targeted therapy.

In this context, we sought to characterize the metabolic reprogramming that takes place during the differentiation/retrodifferentiation of tumor-derived hepatocytes and specify the role of PPARγ in the metabolic adaptation of hepatic CSCs. We used the human HepaRG and HBG-BC2 HCC cell lines, both characterized by high plasticity potential(37,38). For instance, while expression profiles of differentiated-HepaRG cells match with the periportal-type HCC subclass, which discriminates well-differentiated and favorable outcome tumors, gene expression profiles of HepaRG-CSCs are enriched in signatures related to CSCs, metastasis, recurrence and match with Désert’s STEM and Hoshida’s S1 HCC signature, both associated with poor prognosis(3,13). Here, we show that these hepatic CSCs, whose transcriptome matches early stages of mouse development, have reduced mitochondrial biogenesis and cell respiration. In parallel they adopt a glycolytic profile and store lipids into droplets until they leave their slow-proliferating state and commit to a differentiation program. Our results highlight, for the first time, a balanced expression of PPARα and PPARγ along the tumor-derived hepatocyte differentiation/retrodifferentiation process. They reveal a role of PPARγ in the metabolic rewiring of hepatic CSCs, which contributes to chemoresistance. We show that PPARγ has not only antiproliferative but also pro-stemness properties. Our results bring out that PPARγ, whose expression is increased in human HCC STEM subclass, is a negative prognostic factor in three human HCC cohorts. At last, we demonstrate that PPARγ inhibition synergizes with cisplatin or sorafenib, notably by limiting PDK4 expression and reactivating the mitochondrial production of reactive oxygen species (ROS).

## Materials and methods

### Ethics statement, Patient samples and cohorts

Human tissue samples were processed by the Biological Resource Center, Rennes University Hospital (BB-0033-00056) after sample examination by the Anatomic Pathology laboratory. The study protocol complied with French laws and regulations and was approved by INSERM’s Institutional Review Board (number 19-630) in the context of the National Network of Liver Biological Resource Centers. Sample collection was reported to the Ministry of Education and Research (No. DC-2008-338). All the patients provided written informed consent.

Three publicly available HCC transcriptomic datasets were used: Data for the Roessler’s Patient cohort (238 HCCs) are accessible through GEO Series accession number GSE14520; the Cancer Genome Atlas Liver Hepatocellular Carcinoma cohort (TCGA-LIHC; 370 HCCs) through the link https://www.cancer.gov/tcga and the International Cancer Genome Consortium cohort (ICGC; 232 HCCs) project LIRI-JP through the link https://dcc.icgc.org/projects/LIRI-JP.

### Mouse liver development

Two data sets of mouse liver development time course were used: GSE90047 (n=21)(39) and GSE13149 (n=25)(40).

### Cell lines

We used 2 human HCC cell lines established in our laboratory, HBG-BC2(37) (Inserm UMR 1317, ex Inserm U 49) and HepaRG(41) (Inserm UMR 1317, ex Inserm UMR 552, Patent number US7456018). We also used 2 commercial HCC cell lines, HepG2 (ECACC Cat# 85011430, RRID:CVCL_0027), Huh7 (ECACC Cat# 01042712, RRID:CVCL_2957) and Huh-6 cells (RRID:CVCL_4381), a kind gift of Dr. Christine Perret, Institut Cochin(42). HepaRG cells were cultured as previously described(41). Briefly, cells were seeded at 2.7×104 cells/cm^2^ in William’s E medium (Gibco, 22511-022) supplemented with 10% fetal bovine serum (FBS), 100 U/ml penicillin (Gibco, 15070-063), 100 μg/ml streptomycin (Gibco, 15070-063), 5 μg/ml insulin (Sigma-Aldrich, I5500), 5×10^-5^ M hydrocortisone hemisuccinate (Upjohn, Serb) and 2 mM glutamine (Gibco, 25030-024). Progenitors (D4) are cells obtained 4 days after seeding; committed/confluent cells (D15) correspond to cells 2 weeks after seeding. After this stage, the medium was supplemented with 2% DMSO (Sigma-Aldrich, D4540) and the cells cultured for a further 2 weeks to enhance differentiation (differentiated cells, D30) (Fig. S1A). HBG-BC2 cell line was cultured in HepaRG medium without DMSO and maintained 2 weeks at confluency to reach differentiation. HepG2, Huh6 and Huh7 were cultured in DMEM medium (Sigma-Aldrich, M2279) supplemented with 10% FCS, 2 mM glutamine, 100 U/ml penicillin and 100 μg/ml streptomycin. HCC cell spheres and HepaRG-side population were obtained as previously described(11). For spheres, progenitor HepaRG or proliferating HCC cells were cultured in ultra-low attachment plates and stem cell medium consisting of DMEM/F12 medium (Gibco, 11330-032) supplemented with 20% knockout serum replacement (Gibco, 10828-028), 1 mM L-glutamine, 1% nonessential amino acids (Gibco, 11140050), 0.1 mM β-mercaptoethanol (Gibco, 31350010) and 4 ng/ml fibroblast growth factor 2 (Miltenyi Biotec, 130-093-840). HepaRG-SP were obtained from the progenitor population by sorting cells that are able to efflux Hoescht 33342 (11). Experiments on HepaRG-SP cells were performed within 24 hours after seeding in HepaRG medium without DMSO to limit cell proliferation and differentiation. Note that some experiments were performed only with HepaRG-spheres, but not with HepaRG-SP, because of the small amount of SP available.

### Oxygen consumption and glycolysis measurements

Respiration and glycolysis were measured by Seahorse XFe Analyzer (Agilent). Respiration was assessed by successive injections of oligomycin (2 µM), carbonyl cyanide-4 (trifluoromethoxy) phenylhydrazone (FCCP) (1 µM) and the combination of antimycin A/rotenone (1 µM) into the culture medium (Seahorse XF Cell Mito Stress Test Kit, Agilent Technologies, 103015-100). Glutaminolysis was performed by using respiration kit (Seahorse XF Cell Mito Stress Test Kit, Agilent Technologies, 103015-100). 24h before the Seahorse measurements, cells were placed in glutamine-deprived medium or were treated with 1mM of a glutamine antagonist called 6-Diazo-5-oxo-L-norleucine (DON). Glycolysis was assessed by injection of glucose (10 mM), oligomycin (1 µM) and 2-deoxy-D-glucose (2-DG) (50 mM) (Seahorse Glycolysis Stress Test Kit, Agilent Technologies, 103020-100). Analyses were performed with Wave software 2.3.0. Results were normalized to cell number obtained by fluorescence intensity of Hoechst 33342 correlated with nucleus count performed at 460 nm on a POLARstar Omega plate reader (BMG Labtech).

### Assessment of FAO with [U-^14^C]palmitic acid

FAO was assessed by measuring the acid-soluble radiolabeled metabolites resulting from the mitochondrial oxidation of [U-^14^C]palmitic acid as previously described(43). Cells were washed with warm PBS (Gibco, 10010023) and incubated in phenol red-free William’s E medium (Gibco, A1217601) containing 1% fatty acid-free BSA (Sigma-Aldrich, A8806), [U-^14^C]palmitic acid (Perkin Elmer, NEC534050UC), 100 µM cold palmitic acid (Sigma-Aldrich, P5585), 1 mM L-carnitine (Sigma-Aldrich, C0283). After 3 hours of incubation, perchloric acid, final concentration 6%, (Fisher Scientific, 12993564) was added and plates were centrifuged at 2,000g for 10 min. The supernatant was counted for [^14^C]-labeled acid-soluble β-oxidation products using a Tri-Carb 4910TR liquid scintillation counter (PerkinElmer). Results were normalized to cell number as described for oxygen consumption.

### Assessment of De Novo Lipogenesis from ^[2-14C]^acetic acid

*De novo* lipogenesis was assessed by measuring newly synthesized radiolabeled lipids from [2-^14^C]acetic acid, using a protocol from Byrne et al(43,44). Cells were washed with warm PBS and incubated for 3 hours with phenol red-free William’s E medium containing 1% fatty acid-free BSA, [2-^14^C] acetic acid (Perkin Elmer, NEC553050UC) and 50 µM cold acetic acid (Sigma-Aldrich, S5636). Cells were then washed with PBS before adding a mix of hexane/isopropanol (3V/2V) (Sigma-Aldrich, 139386 and I9516) and incubated for 1 hour at room temperature for lipid extraction. After transfer in microtubes, hexane and PBS were added to have a hexane/isopropanol/PBS ratio of 6V/2V/3V. Microtubes were centrifuged at 1000g for 5 min and radiolabeled lipids were counted in the upper phase with a Tri-Carb 4910TR liquid scintillation counter (PerkinElmer). Results were normalized to cell number as described for oxygen consumption.

### Statistical analysis

Microarray data (GSE75752 and GSE112123) were provided by experiments previously performed in the laboratory(11). mRNAs were obtained from biological replicates (n=4) of HepaRG-SP, HepaRG-spheres, and HepaRG cells recovered 4 (HepaRG-progenitor), 15 (HepaRG-committed/confluent) and 30 (HepaRG-differentiated) days of culture (Figure S1A). To determine genes significantly deregulated between progenitors (day 4 after seeding) and differentiated cells (day 30 after seeding), a t-test was performed with the package R Limma. To determine genes significantly deregulated between HepaRG-CSC (spheres and SP) and differentiating (progenitor, committed/confluent and differentiated) cells, a one-way ANOVA was performed. Numerical data comparisons were analyzed using GraphPad Prism software (Version 7.0, GraphPad, San Diego, CA). Significance was assessed by parametric (Student t-test, One-way Anova or Two-way Anova) and nonparametric (Mann-Whitney test or Kuskall-Wallis test) methods according to the results of the normality test. Results are expressed as mean±SEM.

### Data availability statement

The authors confirm that the data supporting the findings of this study are available within the article and/or its Supplementary Materials and methods. Any additional data are available from the corresponding author upon reasonable request.

## Results

### Retrodifferentiation of HepaRG cells alters both the expression of mitochondria-related genes and the mitochondrial network

The acquisition of fetal/hepatoblast characteristics is a well-known feature of hepatic carcinogenesis and is associated with the severity of HCC(4). Retrodifferentiation, which occurs during liver carcinogenesis and HCC progression mirrors foetal development. We previously showed that the transcriptomic programs of SP and Spheres matches those of both the STEM(12,13) / S1(3) HCC subclasses and human embryonic stem cells(11). Here, we sought to clarify to what extent HepaRG differentiation stages (from sphere-forming cells (Spheres), side population (SP), bipotent progenitors, through committed/confluent cells and to differentiated cells, represent different stages of hepatic organogenesis (Fig. S1A). To this end, we integrated HepaRG transcriptomic data with two time-course experiments of mouse liver development (GSE90047(39) and (GSE13149(40)). Hierarchical clustering of transcriptomes from embryonic stages E10.5 through E18.5 and from E11.5 through post-natal days 0 (birth) −21 (weaning) to adult liver, based on the progenitor vs differentiated HepaRG signatures (DEG, P≤0.05, FC>2, Table S1) reveals two clusters (Fig. 1A). Spheres, SP and progenitor cells clustered with early embryonic livers (E10.5 or E11.5 to 14.5) and committed/confluent and differentiated cells clustered with post-natal mouse liver samples (post-natal day 7 to adult liver). These findings indicate that the selected differentiation stages of HepaRG cells correspond to the developmental stages of mouse liver, from the emergence of the liver bud to adult liver and conversely. Therefore, HepaRG cell plasticity may correctly reflect the differentiation/retrodifferentiation processes and the phenotypic diversity of HCCs.

**Figure 1:**
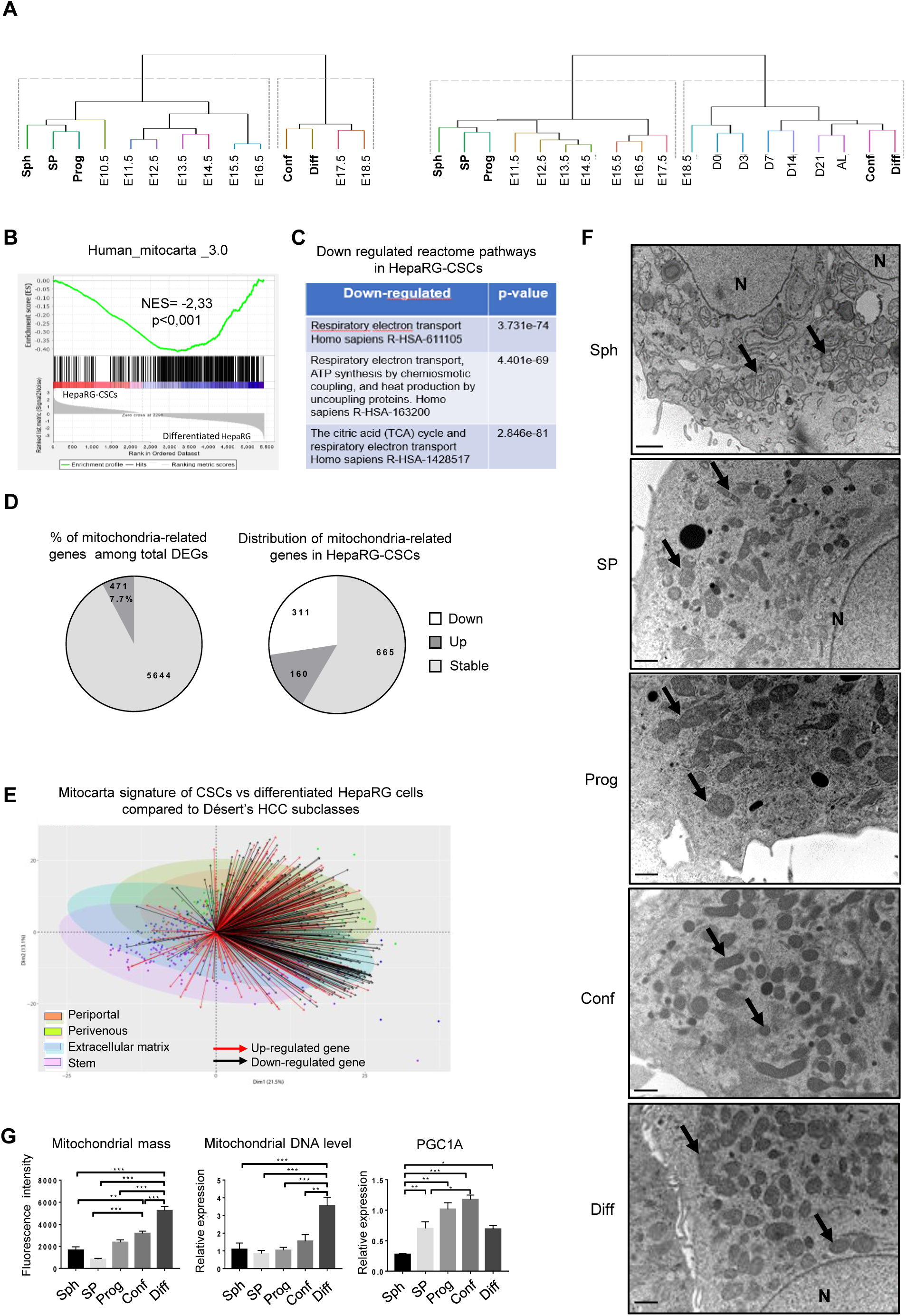
Downregulation of mitochondria-related gene expression in HepaRG-CSCs. (A) Integration of differentially expressed HepaRG genes (DEG, P≤0.05, FC>2) between progenitors (Prog, day 4 after seeding) and differentiated cells (Diff, day 30 after seeding) with orthologs from two mouse liver development mRNA datasets: Left panel: GSE90047(39), from embryonic stages E10.5 through E18.5; Right panel: GSE13149(40), from E11.5 through post-natal days 0 (birth) – 21 (weaning) to adult liver. HepaRG cells: Sph=Spheres; SP=side population; Conf=committed/confluent cells at day 15 after seeding. Mouse liver development: *E,* days post-coitum (detection of the vaginal plug); *D,* post-natal days. Manhattan and ward. D2 were respectively used as distance and clustering method. (B) GSEA plot for mitochondria-related genes referenced in Mitocarta 3.0; NES, normalized enrichment score. (C) Reactome pathways related to DEG in HepaRG-CSCs. (D) Left pie chart: percentage of mitochondria-related genes among total differentially expressed genes (DEG) between immature and differentiating HepaRG groups; Right pie chart: Mitocarta genes upregulated (dark grey) or downregulated (white) in HepaRG-CSCs. (E) Biplot showing the distribution of the 471 DEG belonging to Mitocarta among the Désert’s HCC subclasses: PP=periportal, PV=perivenous, ECM=extracellular matrix. (F) Electron microscopy of HepaRG cells: Sph=spheres, SP=side population, Prog=progenitors, Conf=committed/confluent, Diff=differentiated. Black arrows point out mitochondria; N, nucleus. (G) Mitochondrial DNA assessed by RT-qPCR (n≥4). Mitochondrial mass evaluated by flow cytometry using Mitotracker Green® (n≥6). *PPARGC1A/PGC1A* mRNA expression relative to HepaRG-progenitors (n>=3). *p<0.05, **p<0.01, ***p<0.001.

Then, we characterized the metabolic reprogramming of HepaRG-CSC modelled by HepaRG-spheres and HepaRG-SP(7,37). We identified 6115 differentially expressed genes (DEG, p≤0.05, FC>1.5) between the HepaRG-CSCs and the HepaRG differentiating cells i.e., progenitors, committed/confluent and differentiated cells (Table S2). Unsupervised gene set enrichment analysis (GSEA) reveals that the transcriptomic program of HepaRG-CSCs was negatively correlated with oxidative phosphorylation (OXPHOS), TCA cycle and pyruvate metabolism (Fig. S1B). Accordingly, supervised GSEA shows a negative correlation of HepaRG-CSCs with the Mitocarta gene list(45), a compendium of 1136 human genes encoding mitochondria proteins (Fig. 1B). Among these genes, 471 mitochondria-related genes, accounting for 7.7% of the DEG, are modulated (Fig. 1D, Table S3). Specifically, 160 Mitocarta genes are up-regulated in HepaRG-CSCs while 311 genes, involved in the top three pathways (TCA cycle, mitochondrial respiratory chain and ATP synthesis), are down-regulated (Fig. 1C,D). Next, we integrated the 471 Mitocarta DEG with the 550-gene classifier signature that defines the four HCC subclasses (perivenous, periportal, extracellular matrix, stem) in the Désert’s classification(13). We show that the genes downregulated in HepaRG-CSCs are also down-regulated in the STEM subclass with respect to the periportal and perivenous subclasses (Fig. 1E). This prompted us to study the consequences of transcriptional changes on mitochondria number and network organization. Electron microscopy confirms a reduced number of mitochondria in HepaRG-CSCs compared with differentiated cells (Fig. 1F). This result is supported by both the quantification of mitochondrial DNA content and mitochondrial mass (Fig. 1G). Accordingly, the expression of PPARγ coactivator 1-alpha *(PPARGC1A/PGC1A),* a master regulator of mitochondrial biogenesis, is low in HepaRG-CSCs and gradually increases across the proliferation and epithelial commitment phases of HepaRG-progenitors (Fig. 1G). In addition, confocal microscopy analyses revealed that HepaRG-CSCs harbor a mitochondrial network with significantly fewer branching points (Fig. 2A).

**Figure 2:**
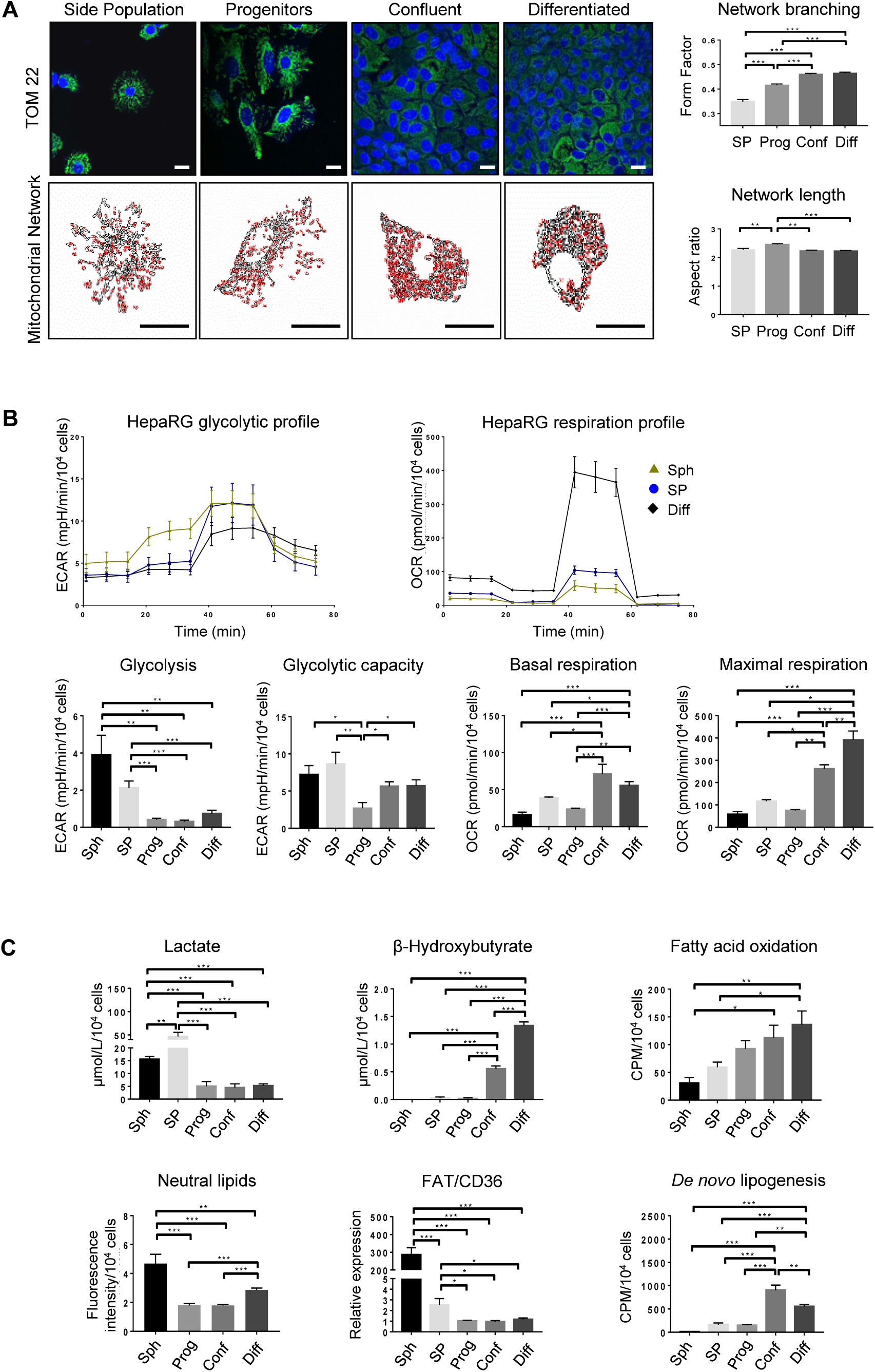
Metabolic reprogramming during differentiation/retrodifferentiation process in HepaRG cells. (A) Confocal microscopy of the mitochondrial protein TOM22 immunostaining in HepaRG cells: SP=side population, Prog=progenitor, Conf=committed/confluent, Diff=differentiated; Bar=20µm. Mitochondrial network length and branching analyzed using ImageJ (n=3); Bars=20µm. (B) Representative glycolytic (extracellular acidification rate, ECAR) and mitochondrial respiration (oxygen consumption rate, OCR) profiles obtained with Seahorse analyzer: Sph=HepaRG-spheres. Glycolysis, glycolytic capacity, basal and maximal respiration (n≥5). (C) Lactate and β-hydroxybutyrate in culture supernatant assessed by absorption spectrophotometry (n≥3). FAO and *de novo* lipogenesis assessed by quantifying the radioactivity subsequent to ^14^C-palmitate and ^14^C-acetate incorporation, respectively (n≥4). FAT/CD36 mRNA expression relative to HepaRG-progenitors (n=4). Neutral lipid content assessed by nile red staining (n≥5). *p<0.05, **p<0.01, ***p<0.001.

### HepaRG-CSCs rewire their metabolism and slow down their proliferation rate

A high-branched mitochondrial network is thought to increase OXPHOS efficiency and energy supply(46). This prompted us to study the metabolism of HepaRG cells during the differentiation/retrodifferentiation process. HepaRG-CSCs adopt a glycolytic profile with both high glycolysis level (Fig. 2B) and increased lactate production (Fig. 2C). Conversely, lower mitochondrial respiration rate is observed in immature than in differentiated HepaRG cells (Fig. 2B). Moreover, FAO (Fig. 2C) and glutaminolysis (Fig. S1C) are low in HepaRG-CSCs. In keeping with the reduced FAO, low β-hydroxybutyrate levels (Fig. 2C) suggests a decrease of ketogenesis. In addition, accumulation of neutral lipids is observed in HepaRG-spheres (Fig. 2C) and -SP(11). This accumulation probably results from increased fatty acids (FAs) uptake through increased expression of the FA transporter FAT/CD36 rather than from *de novo* lipogenesis, which is very low in HepaRG-CSCs (Fig. 2C). It should be noted that HepaRG-spheres and -SP, although both immature cells, display some differential metabolic features. Notably, mitochondrial respiration and FAO are higher in HepaRG-SP. These differences are likely related to the resting state or cell cycle rate of the cells. Indeed, HepaRG-spheres express high levels of early G1 and G1 phase markers (*JUN, CDKN1A, CDK4 and CCND1*) but not the S and M phase markers *CDK1* and *CCNB1*, showing that they are stalled in G1 (Fig. S1D). HepaRG-SP express high levels of early G1 and G1 phase markers (*JUN, FOS, CDK4, CDKN1B*) but also intermediate levels of *CDK1* and *CCNB1*, disclosing a slow-cycling phenotype. As expected, progenitors are active proliferating cells, expressing high levels of S and M phase markers whereas committed/confluent and differentiated HepaRG cells barely express these markers (Fig. S1D).

### HCC cell lines adopt distinct metabolic features according to their plasticity potential and proliferation rates

We completed our study by exploring the metabolism of other HCC cell lines (Fig. S2A). Like HepaRG, HBG-BC2 (BC2) show strong potential for plasticity and differentiation, as evidenced by the inverse expression profile of stemness (CD44) and differentiation (aldolase B) markers in spheres and 15-day-old committed/confluent cells (Fig. 3A). Noteworthy, both HepaRG- and BC2-CSCs form small spheres (Fig. S2A). They have a low *PGC1α* expression (Fig. S2B) and adopt a similar metabolism i.e., a glycolytic profile with high lactate production, low mitochondrial respiration and *de novo* lipogenesis, lipid droplet accumulation and higher *FAT/CD36* expression than their differentiated counterparts (Fig. 3B,C). Of note, FAO and β-hydroxybutyrate levels are very low even in differentiated BC2 cells compared to differentiated HepaRG cells (Fig. 3C). Consistent with the high expression of *FAT/CD36* and lipid accumulation, lipid analysis shows higher amounts of free FAs, triglycerides and cholesterol esters in HepaRG- and BC2-spheres (Fig. 3D). In addition, HepaRG- and BC2-spheres are characterized by higher saturated/unsaturated FA ratio and phospholipid content compared with differentiated cells (Fig. 3D). In contrast to HepaRG and BC2, three other cell lines, Huh6, Huh7 and HepG2, have reduced plasticity and formed spheres that contained proliferative rather than immature cells (Fig. 3A; Fig. S2A). Mitochondrial respiration is high in spheres derived from Huh6, Huh7 and HepG2 cell lines, whereas adherent cells mainly rely on glycolysis (Fig. 3B). In addition, only slight changes in FAO, *FAT/CD36* expression and neutral lipid accumulation are apparent when comparing spheres and adherent proliferative Huh6, Huh7 and HepG2 cells (Fig. 3C).

**Figure 3:**
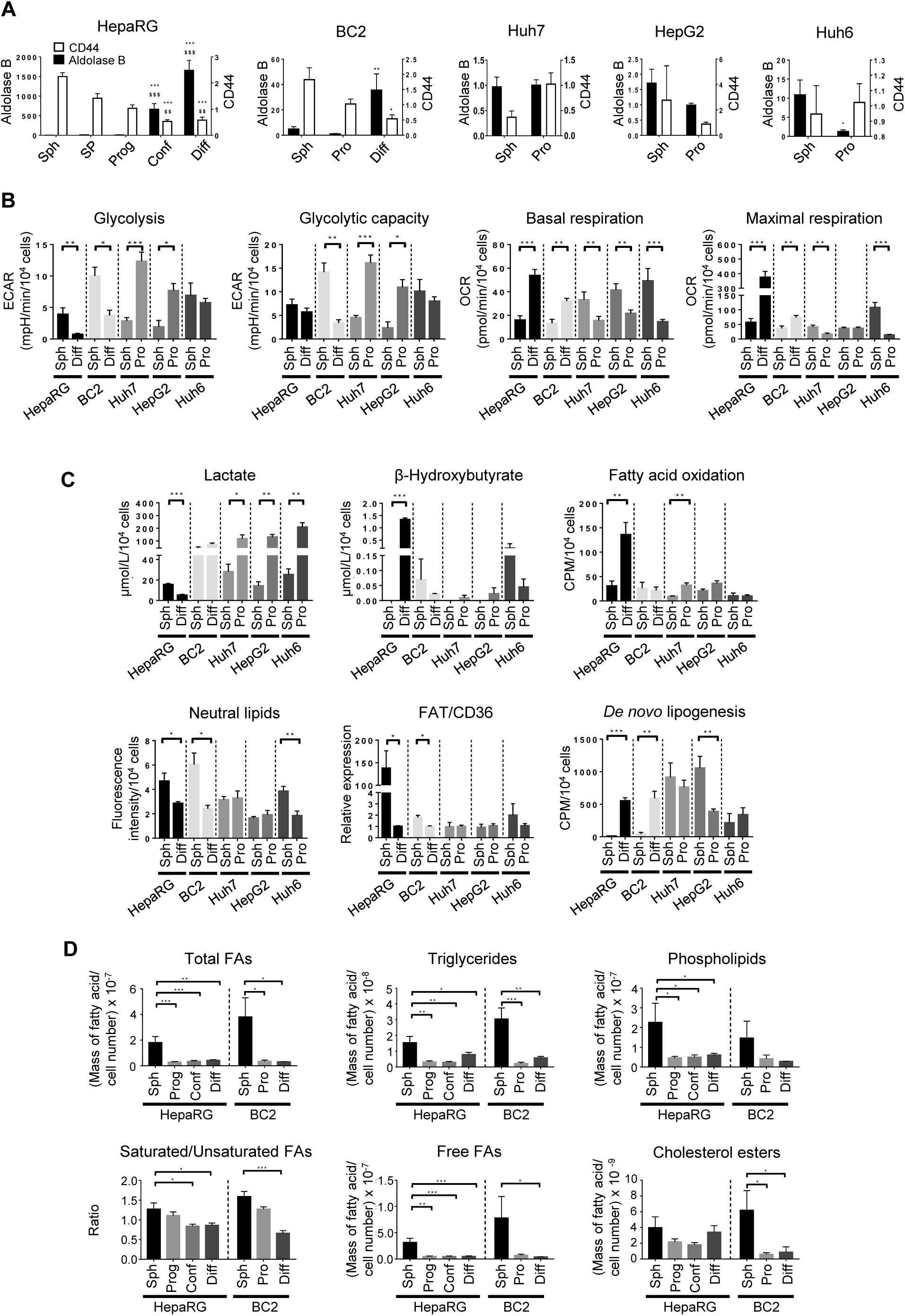
Metabolism rewiring in HCC cell lines. (A) Aldolase B (left y-axis) and CD44 (right y-axis) mRNA expression throughout the culture of HepaRG, BC2, Huh7, HepG2 and Huh6 cell lines: Sph=spheres, SP=side population, Prog=progenitors, Conf=committed/confluent, Diff=differentiated and Pro=proliferative. Results are expressed relative to progenitors for HepaRG or proliferative cells for BC2, Huh7, HepG2 and Huh6 (n≥3). Comparison with spheres *p<0.05, **p<0.01, ***p<0.001; comparison with SP $ p<0.05, $$ p<0.01, $$$ p<0.001 (B) Glycolysis, glycolytic capacity, basal and maximal respiration assessed with Seahorse analyzer for HepaRG, BC2, Huh7, HepG2 and Huh6 (n≥3). (C) Lactate and β-hydroxybutyrate assessed in culture supernatants by absorption spectrophotometry (n≥3). FAO and *de novo* lipogenesis assessed by quantifying the radioactivity subsequent to ^14^C-palmitate and ^14^C-acetate incorporation, respectively (n≥3). FAT/CD36 mRNA expression relative to progenitors for HepaRG and proliferative cells for BC2, Huh7, HepG2 and Huh6 (n>=3). Neutral lipid content assessed by nile red staining (n≥3). (D) Quantification by gas chromatography-mass spectrometry of fatty acids (FA) from cell total lipids (total FAs), triglycerides, phospholipids, cholesterol esters and free FAs in HepaRG and BC2 at different stages of differentiation. The saturated/unsaturated FA ratio is calculated from data obtained with the total lipid pool. Results are normalized by the number of cells (n≥3). *p<0.05, **p<0.01, ***p<0.001.

### High PPARG/PPARA ratio is associated with poor prognosis in human HCC

Changes in lipid metabolism and up-regulated transcription of *PPARG*(11) in HepaRG-CSCs prompted us to further investigate the role of PPAR family members, known as key regulators of cell metabolism. Unlike *PPARA*, *PPARG* expression is high in HepaRG- and BC2-CSCs (Fig. 4A). This likely results from the activation of the PIK3/AKT signaling pathway as Ly294002, which inhibits the AKT upstream activator PI3K, reduces both AKT phosphorylation and *PPARG* expression in HepaRG progenitors (Fig. 4A). Throughout the differentiation process, *PPARG* expression decreases whereas *PPARA* expression changes in the opposite direction, and *PPARD* is fairly stable (Fig. 4A). Interestingly, Kaplan-Meier survival analyses on TCGA-LIHC(47) (Fig. 4B), ICGC and Roessler’s(48) (Fig. S2C,D) HCC patient datasets reveal that high *PPARG* expression is related to worse overall survival. By contrast, expression of *PPARA* is associated with better prognosis in TCGA-LIHC (Fig. 4B) and ICGC cohorts (Fig. S2C,D). Moreover, *PPARG* and *PPARA* are positively and negatively correlated, respectively, with alpha-fetoprotein (AFP), a marker of aggressiveness in HCC, both in TCGA-LIHC and Roessler’s datasets (Fig 4C; Fig. S2E). In addition, high *PPARG* expression matches the poor outcome Désert’s STEM HCC subclass, whereas high *PPARA* expression is found in perivenous-type and periportal-type subclasses, which have better prognosis (Fig. 4D). Fifty-eight HCC tumor tissues were analyzed by immunohistochemistry (IHC), nine of which were positive for PPARγ. Importantly, IHC revealed nuclear localization of PPARγ in the tumor and in tumor nodules invading adjacent stromal tissues (Fig. 4E). Clinical, biological data and risk factors for HCC occurrence are given in Table S4.

**Figure 4:**
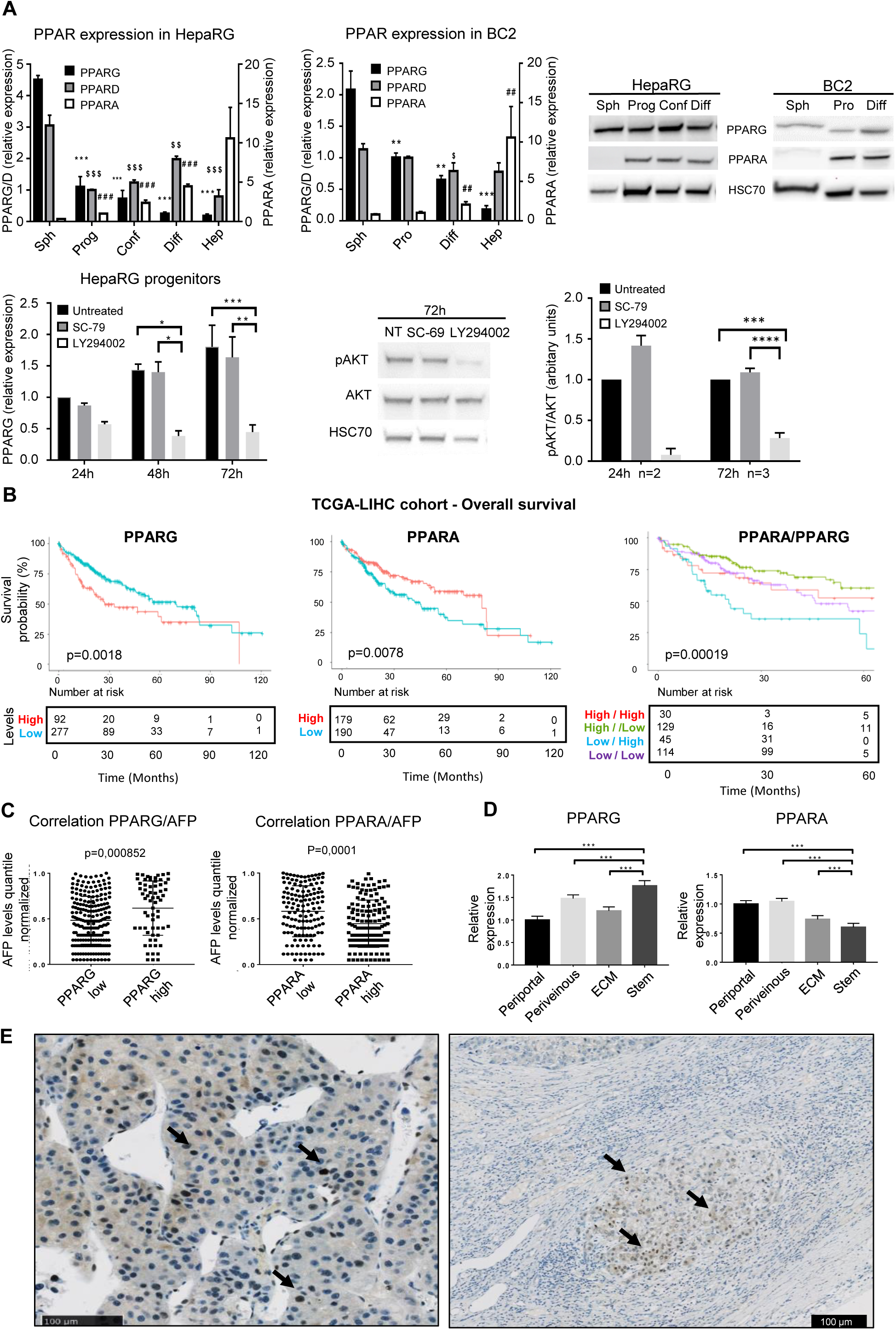
High *PPARG* expression is associated with stem phenotype and poor prognosis in human HCC. (A) Upper left panel: *PPAR* mRNA expression throughout the differentiation process of HepaRG and BC2 cells: Sph=spheres, SP=side population, Prog=progenitors, Conf=committed/confluent, Diff=differentiated, Pro=proliferative, Hep=freshly isolated human hepatocytes. Results are expressed relative to HepaRG-progenitors or BC2-proliferative cells (n≥3). *, $ and # for *PPARG*, *PPARD* and *PPARA*, respectively; $ p<0.05; **, $$, ##p<0,01; ***, $$$, ###p<0,001; all compared with spheres. Upper right panel: Western Blot of PPARγ, PPARα and HSC70 in HepaRG and BC2 cells. Lower left panel: *PPARG* mRNA expression in HepaRG progenitors treated with SC-79 or LY294002 during 24, 48 or 72h. Lower right panel: Western Blot and densitometry of AKT and phosphoAKT (pAKT) in HepaRG progenitors treated with SC-79 or LY294002 during 24 or 72h. *p<0.05; **p<0,01; ***p<0,001. (B) Overall survival according to *PPARG* and/or *PPARA* expression in the TCGA-LIHC cohort. (C) Correlation between *PPARG* and *PPARA* expression and AFP level in the TCGA-LIHC cohort. (D) *PPARG* and *PPARA* expression in the Désert’s HCC subclasses. Results are expressed relative to *PPARG* and *PPARA* levels in the periportal subclass. ***p<0.001. (E) Immunostaining of PPARγ in human HCC. Both the tumor *(left)* and a tumor nodule invading adjacent tissue *(right)* show nuclear and cytoplasmic signal. Black arrows indicate a nuclear localization of PPARγ.

### PPARγ favors stemness and metabolism rewiring in hepatic tumor cells

To get more insight into the role of PPARγ in HCC metabolism, we carried out silencing experiments or agonist treatments using HepaRG- and BC2-spheres, which highly express PPARγ. As expected, the mRNA levels of PPARG is reduced in the presence of siPPARG) whereas PPARγ-agonist rosiglitazone does not affect its expression (Fig 5A and Fig. S3A). Inhibiting PPARγ expression by siRNA transfection drastically reduces the sphere number and causes significant cell death whereas treatment with the PPARγ agonist rosiglitazone induces no change in cell viability (Fig. 5A). Due to cell number requirement, we next used proliferative HepaRG-progenitors and BC2 cells that also express PPARγ at significant levels (Fig. 5B-E, Fig. 6A-C). Treatment with rosiglitazone slows down cell proliferation, thus leading to empty areas in culture plates and reduces ATP levels unrelated to apoptosis induction (Fig. 5B). Treatment also increases the expression of known PPARγ target genes such as *PDK4*, *FAT/CD36* and perilipin2 (*PLIN2*), which contributes to the formation and stability of lipid droplets (Fig. 5C). Interestingly, rosiglitazone treatment results in decreased expression of the epithelial marker E-cadherin (*CDH1*) associated with increased expression of the transcription factor *SNAIL* and the stem cell marker *KLF4*, showing that cells commit to EMT and stemness (Fig. 5C). Consistent with rosiglitazone-induced retrodifferentiation, cells have a less extensive and branched mitochondrial network (Fig. 5D; Fig. S4B) as well as lower mitochondrial respiration (Fig. 5E). In accordance, attenuation of PPARγ expression (RNA and protein levels) by siRNA (Fig. 6B; Fig. S3B) induces a decreased expression of *PDK4* and *PLIN2* (Fig. 6B). This decrease is associated with increased respiratory capacities and proton leak reflecting suboptimal efficacy of the respiratory chain (Fig. 6C). In addition, *PPARG*-invalidated cells contain an increased level of mitochondrial DNA and exhibit an extensive and high-branched mitochondrial network at 48h (Fig. 5D) and 72h (Fig. S4B). Overall, *PPARG* invalidation leads to decreased cell viability (Fig. 6A and Fig. 7C).

**Figure 5:**
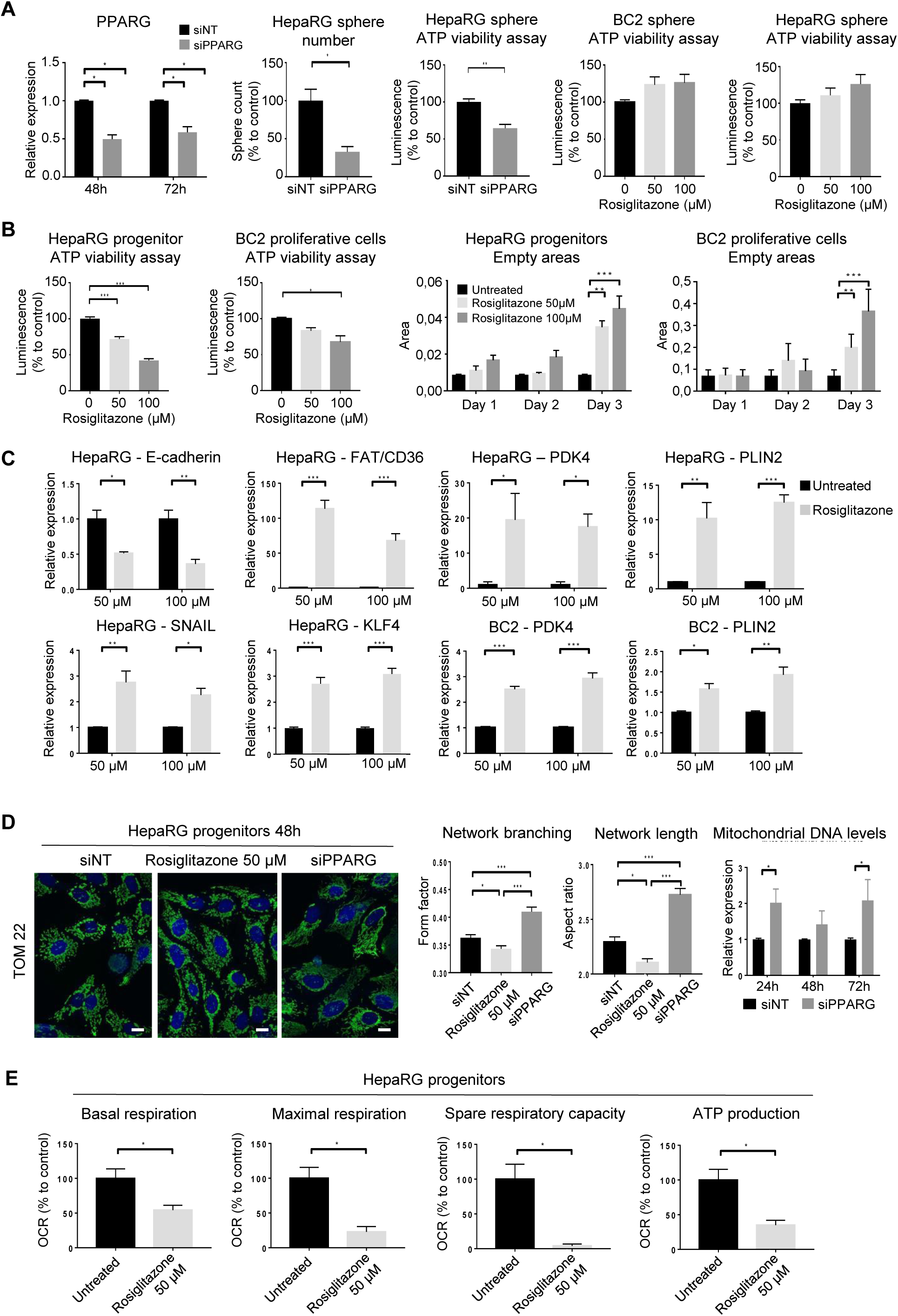
PPARγ activation triggers retrodifferentiation and metabolic reprogramming. (A) *PPARG* mRNA expression 48h after transfection of HepaRG-spheres with siNon-targeting (siNT) and siPPARG (n=3). Sphere number and ATP viability assay 48h after seeding of transfected HepaRG cells in sphere culture medium (n=3). ATP viability assay of HepaRG- and -spheres after treatment with 50µM or 100µM rosiglitazone during 72h (n=4). (B) ATP viability assay of HepaRG-progenitors and BC2-proliferative cells after treatment with 50µM or 100µM rosiglitazone during 72h (n=3). HepaRG and BC2 proliferation assessed by quantifying empty areas using ImageJ, three days after seeding. (n=3). (C) mRNA expression of E-cadherin (*CDH1*), fatty acid transporter *FAT/CD36*, *PDK4*, perilipin2 (*PLIN2*), EMT-inducing transcription factor *SNAIL* and stem-related marker *KLF4* in HepaRG-progenitors and BC2- proliferative cells after treatment with rosiglitazone (50 or 100µM) during 72h (n=3). (D) Confocal microscopy of mitochondrial protein TOM22 immunostaining in HepaRG-progenitors 48h after treatment with 50µM rosiglitazone or transfection with siPPARG. Bar=10µm. Mitochondrial network branching and length analyzed using ImageJ (n=3). Mitochondrial DNA assessed by RT-qPCR, 24h, 48h and 72h after transfection of HepaRG-progenitors with siPPARG (n=4). (E) Basal respiration, maximal respiration, spare respiratory capacity and respiration linked to ATP production assessed with Seahorse analyzer in HepaRG-progenitors treated by rosiglitazone 50µM during 48h (n=3). Results are expressed relative to untreated cells or siNT. *p<0.05, **p<0.01, ***p<0.001

**Figure 6:**
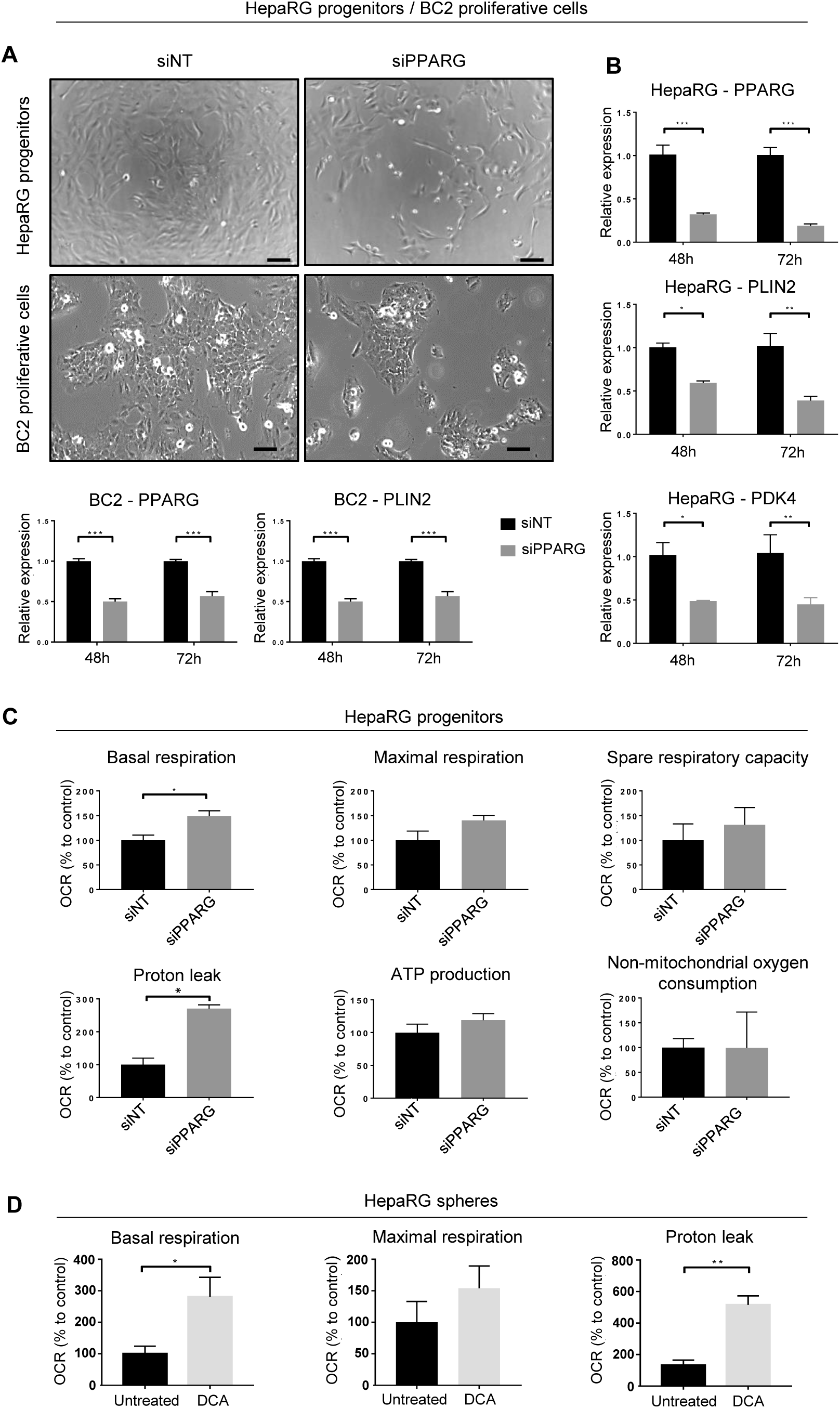
PPARγ inhibition reduces PDK4 and PLIN2 expression and restores respiration. (A) Phase-contrast microscopy of HepaRG-progenitors and BC2-proliferative cells 48h after transfection with siPPARG. Bar=100µm. (B) mRNA expression of *PPARG*, *PLIN2* and *PDK4*, 48h and 72h after transfection of siPPARG in HepaRG-progenitors and BC2-proliferative cells (n=3). (C) Basal respiration, maximal respiration, spare respiratory capacity, proton leak, respiration linked to ATP production and non-mitochondrial oxygen consumption, assessed with Seahorse analyzer in HepaRG-progenitors, 48h after transfection of siPPARG (n=3). Results are expressed relative to siNT. (D) Basal respiration, maximal respiration and proton leak assessed with Seahorse analyzer in HepaRG-spheres treated by DCA (50mM) during 4h. Results are expressed relative to untreated HepaRG-spheres (n>=3). *p<0.05, **p<0.01, ***p<0.001.

**Figure 7:**
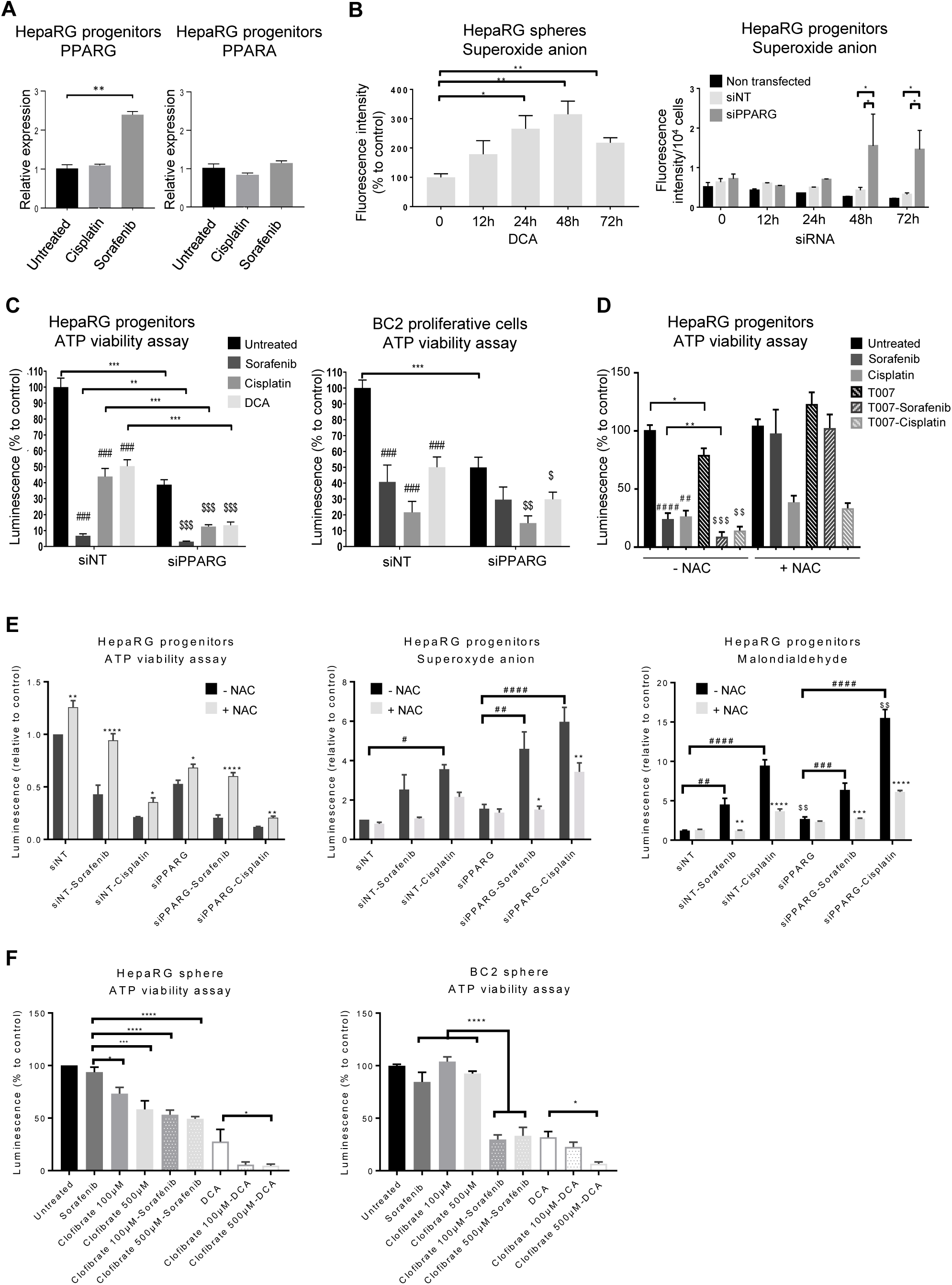
PPARγ inhibition or agonist-mediated activation of PPARα improves the efficacy of standard HCC chemotherapies in vitro. (A) *PPARG* and *PPARA* expression in HepaRG progenitors after treatment during 72h with cisplatin (10µl/ml) or sorafenib (2.5µM) (n=4). **p<0,01 (B) Superoxide anion production (ROS) assessed by Mitosox® in HepaRG-spheres after 50 µM dicloroacetate (DCA) treatment during 12h, 24h, 48h and 72h (n=4) or in HepaRG progenitors after 12h, 24h, 48h and 72h transfection with siPPARG or siNT (n=4). *p<0.05; **p<0,01 (C) Viability of HepaRG- progenitors and BC2-proliferative cells transfected with siPPARG or siNT and treated with cisplatin (10µg/ml for HepaRG; 20µg/ml for BC2) or sorafenib (2.5µM for HepaRG; 15µM for BC2) or 50mM DCA during 48h. Results are expressed relative to untransfected/untreated cells (n=4 for HepaRG and n=3 for BC2). *, #, $ in comparison to siNT, untreated siNT and untreated siPPARG, respectively. (D) Viability of HepaRG-progenitors pre-incubated 24h or not with N-acetyl-cysteine (NAC, 1mM) and treated during 48h with cisplatin (10µg/ml) or sorafenib (2.5µM) combined or not with T0070907 (10µM). Results are expressed relative to untreated cells (n=3). *, T0070907 vs untreated; #, cisplatin or sorafenib vs untreated; $, cisplatin or sorafenib vs cisplatin or sorafenib combined with T0070907. (E) Viability, superoxide anion production, lipid peroxidation assessed by malondialdehyde formation in HepaRG transfected with siPPARG or siNT, pre-incubated 48h or not with NAC (1mM) and treated during 48h with cisplatin (10µg/ml) or sorafenib (2.5µM). *, NAC vs without NAC; #, cisplatin or sorafenib vs untreated; $, siPPARG vs siNT. (F) Viability of HepaRG- and BC2-spheres pre-treated 24h with clofibrate (100 or 500µM) and then treated with sorafenib (2.5µM for HepaRG; 15µM for BC2) and/or DCA (50mM) during 72h. Results are expressed relative to untreated cells (n=4 for HepaRG and n=3 for BC2). *, #, $ p<0.05; **, $$ p<0.01; ***, ###, $$$ p<0.001. ****, #### p<0.0001.

### PPAR targeting improves efficacy of HCC therapies *in vitro*

We have previously shown that HepaRG-progenitors resistant to sorafenib or cisplatin express increased mRNA levels of *PDK4* and that the combination of chemotherapy with the PDK4 inhibitor (DCA) is effective in killing immature HCC cells(11). Here, we show that sorafenib-resistant progenitors also express higher levels of *PPARG*, a master regulator of *PDK4,* while their *PPARA* expression remains unchanged (Fig 7A). Interestingly, targeting either PDK4 with DCA or *PPARG* with siRNA reactivates mitochondrial respiration (Fig. 6C-D) and increases ROS production (peroxide and superoxide anions) (Fig. 7B and Fig. S3C), which hepatic CSCs fail to scavenge, unlike differentiated cells (Fig. S3C). Indeed, most of the metabolic redox regulatory genes are less expressed in immature than in differentiated HepaRG cells (Fig. S4A). As consequences, similar to DCA treatment, si*PPARG* transfection induces cell death of HepaRG-progenitors and BC2 proliferating cells and the dual inhibition of PPARγ by siRNA transfection and PDK4 by DCA proves synergistic in reducing cell viability (Fig. 7C). In addition, *PPARG* invalidation or PPARγ antagonist T0070907 also synergizes with sorafenib or cisplastin (Fig. 7C,D). Decrease in cell viability is associated with increased ROS production and lipid peroxidation as shown by malondialdehyde accumulation in culture medium (Fig. 7E). To confirm the involvement of ROS in cell death mediated by PPARG inhibition, we aimed to scavenge ROS by N-acetylcysteine (NAC). As expected, NAC reduces ROS production, lipid peroxidation and cell death (Fig. 7E).

The better outcome of patients with high *PPARA* expression in the TCGA-LIHC and ICGC cohorts could result from the metabolic effects of PPARα (Fig. 4B and Fig. S2C,D). We hypothesized that, unlike PPARγ, PPARα activation increases the mitochondrial activity without generating high levels of PDK4, which contributes to chemotherapy resistance in hepatic CSCs. Accordingly, the PPARα agonist clofibrate slightly increases mitochondrial network length in HepaRG-progenitors (Fig. S4C), slightly modulates *PPARG* expression and induces PDK4, but at much lower levels than PPARγ agonist rosiglitazone (Fig. S3A). Last, we confirmed this hypothesis using HepaRG- and BC2-spheres. Treatment with clofibrate alone does not reduce cell viability, but the combination of clofibrate with sorafenib, cisplatin or DCA reduces cell viability more than monotherapies (Fig. 7F).

## Discussion

Using several HCC cell lines, including HepaRG and HBG-BC2 cells that exhibit well-characterized differentiation and retrodifferentiation potential(7,11,38), we showed that hepatic CSCs derived from retrodifferentiation of differentiated tumor-derived hepatocytes express *PPARG* and adopt a specific metabolic profile associated with chemoresistance. Although oxidative metabolism is critical for differentiated tumor-derived hepatocytes, hepatic CSCs are characterized by low OXPHOS, mitochondrial FAO, *de novo* lipogenesis and high glycolytic activity. Hepatic CSCs also accumulate lipid droplets, where the triglycerides can be mobilized to fulfill energy needs. Importantly, this metabolic phenotype is reversible and not mutation-driven. Indeed, HepaRG cells are mutated in the *hTERT* promoter(49), whereas BC2 cells are mutated in both *TP53* (H214R substitution associated with second allele loss)(38) and *CTNNB1* (S45A substitution associated with second allele loss, Cavard C., personal communication).

Metabolic reprogramming gives hepatic cancer cells the ability to adapt to energy requirements and environmental constraints. To avoid oxidative stress, cancer cells can either develop a proficient ROS scavenging system or undergo metabolic reprogramming to curb ROS production. Notably, a glycolytic phenotype may allow cells to produce enough energy to maintain their functions while reducing ROS production and oxidative damages(50–52). In this work, we show that hepatic CSCs exhibit reduced mitochondrial biogenesis and harbor a mitochondrial network with fewer branching points than differentiated tumor-derived hepatocytes. Such a fragmented network, which we have shown to be linked to PPARG expression in HepaRG-CSCs, would favor the adoption of a glycolytic profile and self-renewal properties of stem cells rather than commitment in cell differentiation(53). Unlike stem cells that produce antioxidants and/or express antioxidant enzymes (51), HepaRG-CSCs express low level of anti-oxidant enzymes(11). As a result, recovery of mitochondrial respiration by silencing *PPARG* and/or inhibiting its target gene *PDK4* leads to increased ROS production and cell death in both HepaRG- and BC2 spheres. Indeed, PDK4-induced metabolic shift is beneficial for CSCs because it limits the production of mitochondrial ROS that can oxidatively damage lipids, proteins and nucleic acids, especially when cellular antioxidant defenses are overwhelmed(52). In addition, PDK4 could restrain ferroptosis by limiting iron-dependent phospholipid peroxidation that mainly occurs when the TCA cycle is functional(54). Therefore, glycolytic reprogramming and high expression of PPARG/PDK4 in retrodifferentiated-derived hepatic CSCs contribute to limit ROS production and the resulting damages.

Besides a glycolytic phenotype, hepatic CSCs also exhibit an altered lipid metabolism that may help prevent oxidative damages. Hepatic CSCs store more saturated than unsaturated lipids, the latter being prone to peroxidative attack(55). Thus, high levels of saturated FAs are thought to decrease lipid peroxidation and subsequent ROS-induced damages by diluting polyunsaturated FAs in cell membranes(56). Both low *de novo* lipogenesis and desaturase activity probably explain the low level of unsaturated lipids in hepatic CSCs. Low *de novo* lipogenesis might also force hepatic CSCs to use extracellular FAs, in particular through increased expression of the membrane transporter FAT/CD36. Once taken up, lipids are not shuttled into mitochondria to produce energy, but rather stored in droplets. In consistency with these observations, accumulation of lipid droplets has been associated to stemness in various cancer types(57–59). By sequestrating lipids, the droplet core provides a protective environment limiting lipid oxidation and oxidative stress(60). In addition, lipid storage likely provides a source of energy that can be quickly mobilized to meet energy requirements during cell proliferation or differentiation. Accordingly, decreased lipid content is concomitant with restoration of FAO in proliferating and differentiating HepaRG cells. Furthermore, HepaRG-spheres are arrested in G1 phase of the cell cycle and harbor more lipid droplets than HepaRG-SP(11), which express higher level of the S and M phase markers. Interestingly, the phenotype of HepaRG-SP displays similarities with hepatic tumor-initiating stem-like cells described by Chen et al, which have self-renewal ability, low OXPHOS but active FAO(50). Of note, the pluripotency transcription factor NANOG that represses OXPHOS and mitochondrial ROS production in these hepatic tumor-initiating cells is expressed in HepaRG-CSCs(7,11). In addition, HepaRG- and BC2- CSCs express the transcription factor *KLF4*, which drives stemness phenotype during somatic cell reprogramming. KLF4 also favors EPCAM and CD133 expression when overexpressed in Huh7 cells(61). Thus, the retrodifferentiation process leads differentiated malignant hepatocytes to adopt a CSC-like phenotype characterized by low proliferation rate, energy metabolism limiting ROS production and lipid storage for future needs.

The nuclear receptors PPARs orchestrate lipid and glucose metabolism and emerging evidence indicates a complex relationship between PPARs and HCC development. Here, we demonstrate that the differentiation/retrodifferentiation process in HCC cells is modulated by a balanced expression of PPARα/γ isoforms. PPARα, the main isoform expressed in the liver, is found in differentiated tumor-derived hepatocytes, associated with high PGC1α expression, branched mitochondrial network and OXPHOS. High *PPARα* expression is also associated with favorable outcome in HCCs in two patient datasets (TCGA-LIHC and ICGC). However, the cellular localization of PPARα should be considered with more attention for the stratification within the cohorts because patients with reduced nuclear localization of PPARα have a poorer prognosis(22). On the other hand, PPARγ, which is not expressed in the basal state in differentiated hepatocytes, is found in steatosis(21) and steatosis-associated liver cancers(27,29). Unexpectedly, we found a PPARγ expression in hepatic CSCs. This could result from high AKT signaling and low expression of HNFα1, a crucial transcription factor for hepatocyte differentiation we previously reported in HepaRG-spheres and -SP(11). Indeed, it was recently shown that AKT2 phosphorylation promotes PPARγ expression through negative regulation of HNF1α(27). Unlike PPARα, high level of PPARγ is associated with low level of PGC1α, fragmented mitochondrial network and negative regulation of numerous genes involved in the TCA cycle and OXPHOS. In line with these observations, the balance *PPARA/PPARG* affects the prognosis of HCCs and high *PPARG* expression is associated with poor overall survival in three independent HCC cohorts (TCGA-LIHC, ICGC and Roessler), totaling 843 patients. Yu et al previously demonstrated that PPARγ limits HCC cell proliferation and growth(26). Consistent with this data, our results show that PPARγ activation by the agonist rosiglitazone reduces hepatic CSC proliferation. Importantly, PPARγ-induced metabolic rewiring, associated with acquisition of stemness features, leads to chemoresistance of HCC cells. These observations are discrepant with a recent work showing that PPARγ is required for PGC1α-induced inhibition of Wnt/βcatenin/PDK1 axis and metastasis in HCCs(25). This discrepancy might be related to the low expression of PGC1α in hepatic CSCs and/or to the ratio PPARα/PPARγ, which may be different depending on the differentiation state of the cells studied. Interestingly, in line with the deleterious role of PPARγ in HCC, Xiong et al. recently showed that PPARγ signaling promotes resistance to checkpoint inhibitors via induction of VEGFA transcription, which drives an immunosuppressive environment with myeloid-derived suppressor cell expansion and CD8 T cell dysfunction (62). In agreement with this observation, we previously observed that HepaRG-CSC expressed VEGFA(11). In summary, our work provides new insights into PPARγ activation in HCC cells. PPARγ 1) induces the expression of the stem cell marker, KLF4, 2) modulates the expression of the EMT markers SNAIL and E-Cadherin, 3) increases the expression of the FA transporter FAT/CD36 and PLIN2, a lipid droplet-stabilizing protein, 4) enhances the expression of PDK4, a pivotal actor of metabolic flexibility and 5) alters the mitochondrial network and respiration capacity. In conclusion, we demonstrate that the balance between PPAR isoforms modulates the metabolic and phenotypic reprogramming in HCC cells. Specifically, PPARγ rewires cell metabolism toward that of stem cells, contributing to chemoresistance. Our work brings out an underestimated function of PPARγ in the process of liver cell differentiation/retrodifferentiation and strengthens the interest of evaluating cell differentiation modulators as new therapeutic options for HCC.

## Abbreviations used in this paper

CSCs: cancer stem cells
DCA: dichloroacetate
EMT: epithelial-to-mesenchymal transition
FAO: fatty acid oxidation
HCCs: hepatocellular carcinomas
HNF4: hepatocyte nuclear factor-4
OXPHOS: oxidative phosphorylation
NAC: N-Acetyl-Cysteine
PDK4: pyruvate dehydrogenase kinase 4
TCA: tricarboxylic acid
PPAR: peroxisome proliferator-activated receptor
ROS: Reactive oxygen species
SP: side population

## Authors’ contributions

A.C., F.C. conceived the study, supervised the work and analyzed the data. Y.D., L.M., S.P., C.R., K.F., L.C., L.D., D.C., A.B., C.S., C.R. performed the experiments and analyzed the data. C.A., C.B., B.C., CA, V.R., O.M. and B.F. analyzed the data. Y.D., F.C. and A.C. wrote the manuscript.

## Acknowledgements

The authors thank Pr. Béatrice Desvergne, Faculty of Biology and Medicine, Center for Integrative Genomics, University of Lausanne, for critical reading of the manuscript; Dr Guilia. Bertolin for her advice during the mitochondrial network analysis; Drs. Mireille Desille and Bruno Turlin, CRB-Santé, UAR Biosit, Biogenouest, Rennes, France for tissue bank management; Pascale Bellaud (H2P2 platform, University Rennes 1, Rennes, France); Isabelle Cannie and Tifenn Le Charpentier for technical support; Tyéfenn Bouvron, for pre-processing of publicly available raw RNAseq data from mouse liver development models and Patricia Jouas, Adina Pascu and Thomas Poussou for secretarial and informatics support. The authors also thank the “Génomique Santé”, “ImPACcell”, “flow cytometry”, “H2P2” Core facilities from Biogenouest, UAR Biosit, University of Rennes1.

