## Supplemental information and figures for "PPARγ, a key modulator of metabolic reprogramming, stemness and chemoresistance associated with retrodifferentiation in human hepatocellular carcinomas"

#### Supplementary Figure 1

(A) Phase-contrast microscopy of HepaRG cells during differentiation/retrodifferentiation. Bar=50μm. (B) Non-supervised gene set enrichment analysis (GSEA) plot from KEGG gene set database performed using the differentially expressed genes (DEG) between immature

(Sphere and SP) and differentiating (progenitors, committed/confluent and differentiated) HepaRG cell groups. (C) Basal and maximal respiration assessed with Seahorse® analyzer. HepaRG were treated 24h with a glutamine antagonist (DON) or cultured in medium without glutamine (n=3). \* p<0.05, \*\* p<0.01, \*\*\* p<0.001 in comparison with the sphere condition; \$ p<0.05, \$\$ p<0.01, \$\$\$ p<0.001 in comparison with the progenitor condition. (D) mRNA expression of *JUN*, *FOS*, *CDKN1A*, *CDKN1B*, *CDK4*, *CCND1*, *CDK1* and *CCNB1*. Results are expressed as relative to HepaRG progenitors, arbitrary set to 1 (n=4). \* p<0.05, \*\* p<0.01, \*\*\* p<0.001.

##### Supplementary Figure 2

Phase-contrast microscopy of BC2, Huh7, HepG2 and Huh6 in sphere conformation (upper panel) and proliferative or differentiated stages (lower panel); Bar=50µm. (B) mRNA expression of *PPARGC1A*/*PGC1a* in HepaRG, BC2, Huh-7, HepG2 and Huh6 cells at proliferative/differentiated stages or in sphere conformation. Results are expressed as relative to differentiated cells for HepaRG and BC2 cell lines or relative to proliferative cells for Huh7, HepG2 and Huh6 cell lines, arbitrary set to 1 (n = 3). \* p<0.05, \*\* p<0.01, \*\*\* p<0.001. (C) Overall survivals according *PPARA*/*PPARG* expression in the ICGC and Roessler cohorts. (D) Correlation between *PPARA*/*PPARG* expression and AFP levels in the Roessler cohort.

##### Supplementary Figure 3

(A) mRNA expression of *PPARG*, *PPARA* and *PDK4* in HepaRG progenitors after treatment with increasing concentrations of rosiglitazone (50 and 100µM) or clofibrate (50, 100, 300, 500 and 1000µM) during 24 or 48h. Results are expressed as relative to the untreated cells, arbitrary set to 1 (n=3). \*p<0.05, \*\*\* p<0.001. (B) Left panel: *PPARγ* expression assessed by western blot 24, 48 or 72h after transfection of HepaRG progenitors by 4 different si*PPARG*; Right

panel: PPARγ expression assessed by western blot 72h after transfection of HepaRG progenitors by siPPARG-4 (n=3). HSC70 protein was used as normalization protein. (C) Upper left panel: production of peroxides (reactive oxygen species) by HepaRG-spheres after DCA treatment during 12h, 24h, 48h and 72h, assessed using H2DCFDA (n=3); Upper right panel: production of peroxides by HepaRG-progenitors after siRNA transfection during 12h, 24h, 48h and 72h, assessed using H2DCFDA. \*p<0.05, \*\* p<0.01, \*\*\* p<0.001. Lower left panel: Phase-contrast microscopy of differentiated HepaRG cells; Lower right panel: production of peroxide and superoxide anions (reactive oxygen species) by differentiated HepaRG cells after DCA treatment during 12h, 24h, 48h and 72h, assessed using H2DCFDA or Mitosox®, respectively (n=3). \*\* p<0.01.

###### **Supplementary Figure 4**

(A) Heatmap of metabolic redox regulatory gene among the 6115 differentially expressed genes (DEG, $p \leq 0.05$, $FC > 1.5$) identified between the HepaRG-CSCs (sphere and side population) and the HepaRG differentiating cells i.e., progenitors, committed/confluent and differentiated cells. (B) Immunostaining of the mitochondrial protein TOM22 in HepaRG progenitors 72h after treatment with 50μM rosiglitazone or transfection by siPPARG. Image acquisition was performed by confocal microscopy. Bar=10μm. Mitochondrial network branching and length were analyzed using ImageJ software (n=3). \*\*\*p<0.001. (C) Immunostaining of the mitochondrial protein TOM22 in HepaRG progenitors 48h after treatment with 100μM clofibrate. Image acquisition was performed by confocal microscopy. Bar=10μm. Mitochondrial network branching and length were analyzed using ImageJ software (n=3). \* p<0.05.

A

#### HepaRG

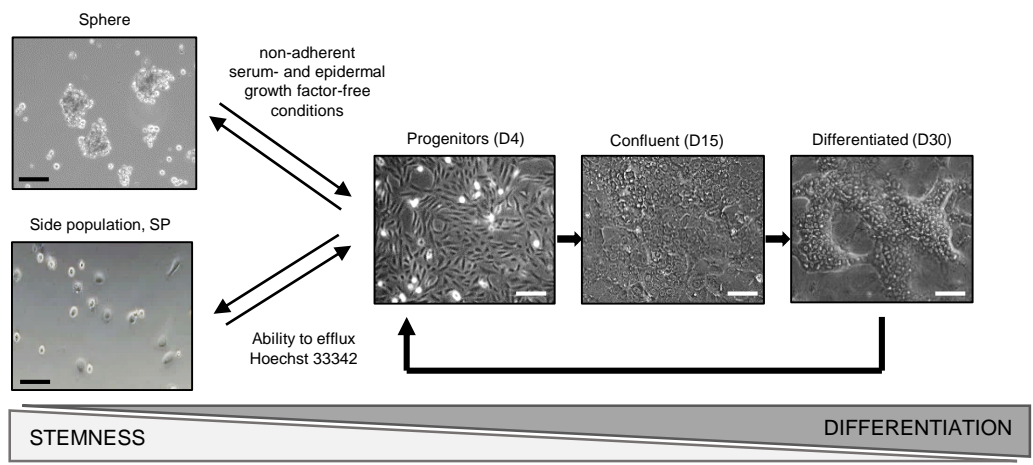

B

KEGG\_OXIDATIVE\_PHOSPHORYLATION
HepaRG stem vs differentiated

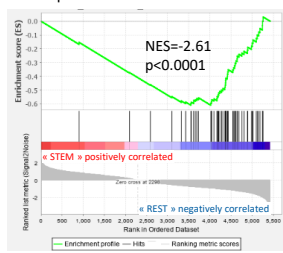

KEGG\_CITRATE\_CYCLE\_TCA\_CYCLE
HepaRG stem vs differentiated

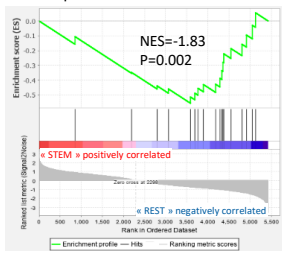

KEGG\_PYRUVATE\_METABOLISM
HepaRG stem vs differentiated

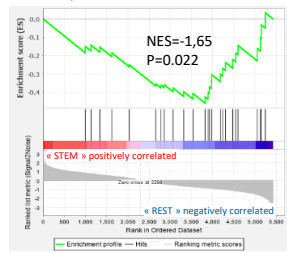

C

Basal respiration

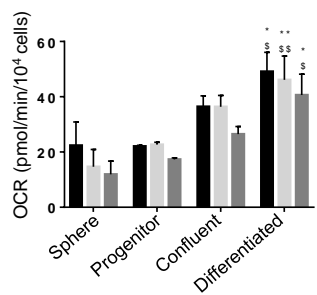

Maximal respiration

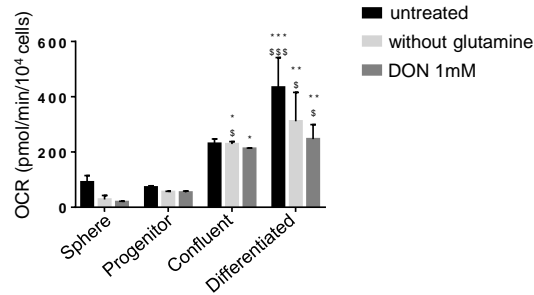

D

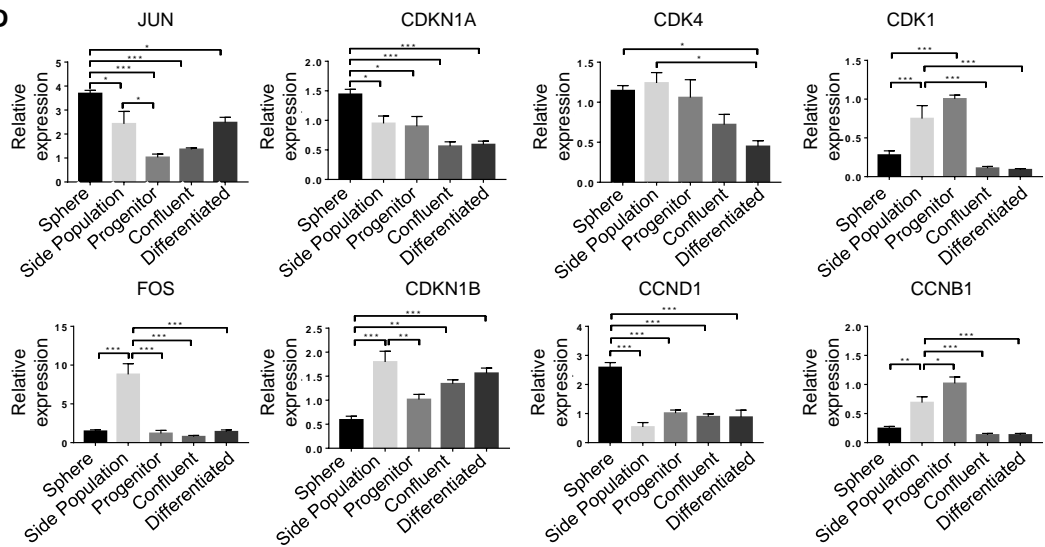

Supplementary Figure 1

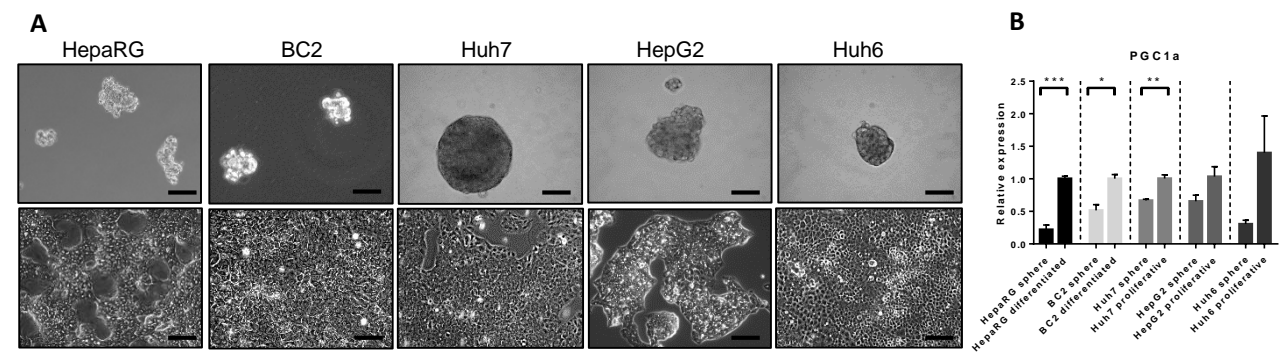

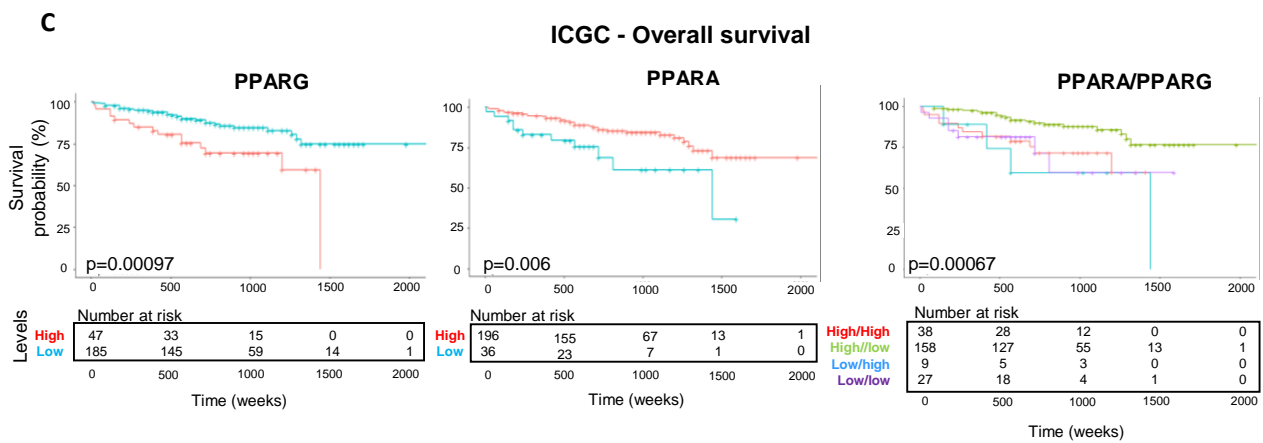

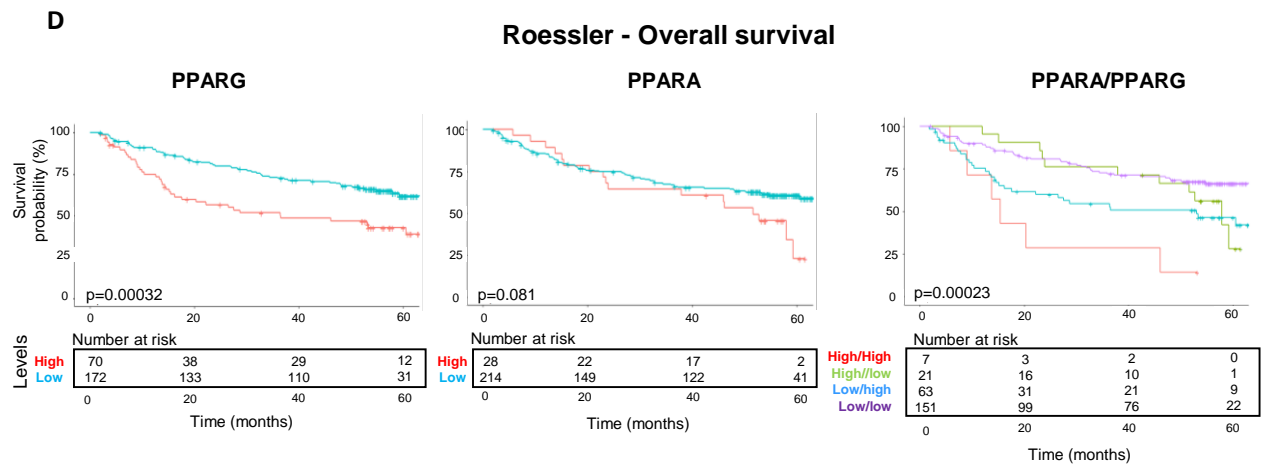

**E**

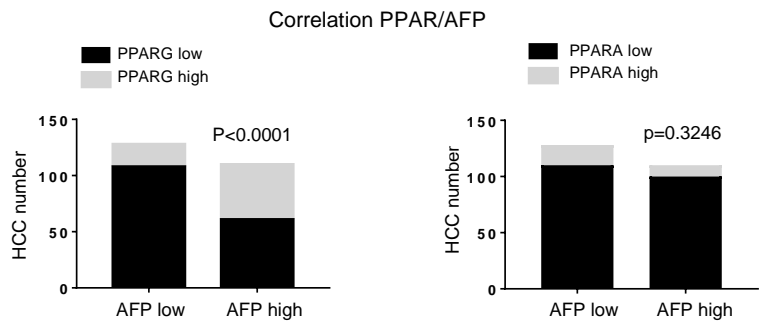

Supplementary Figure 2

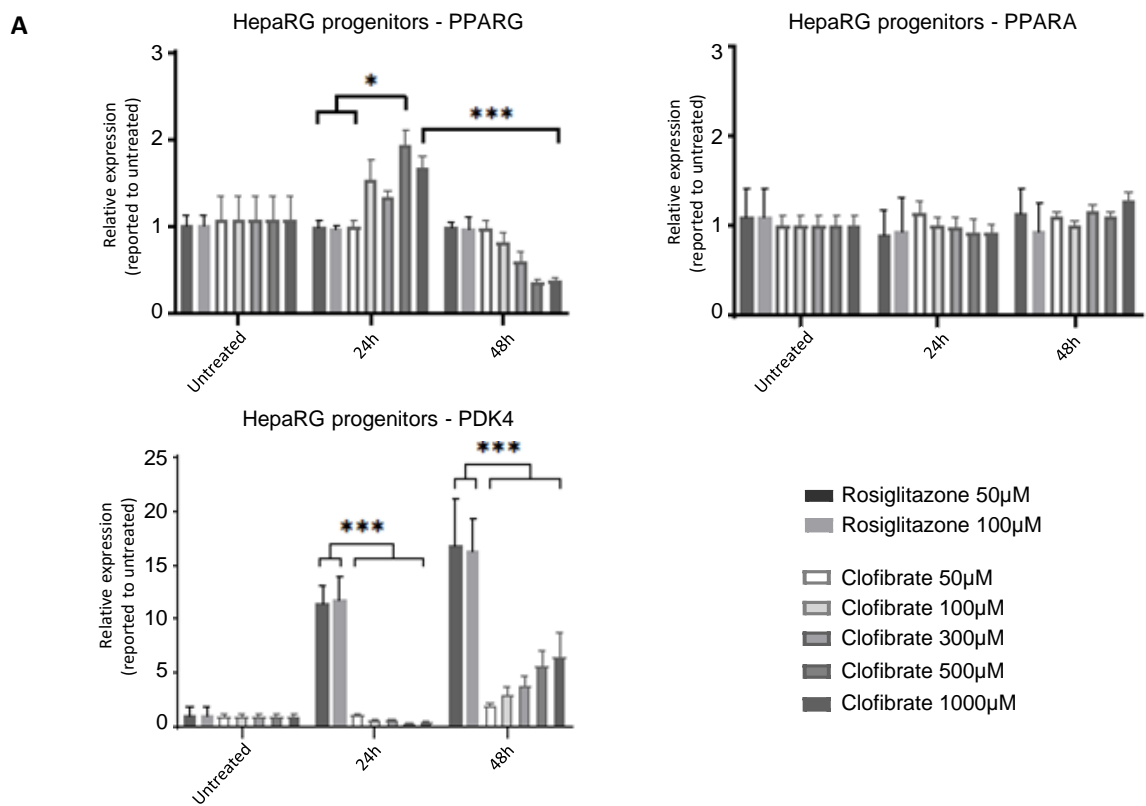

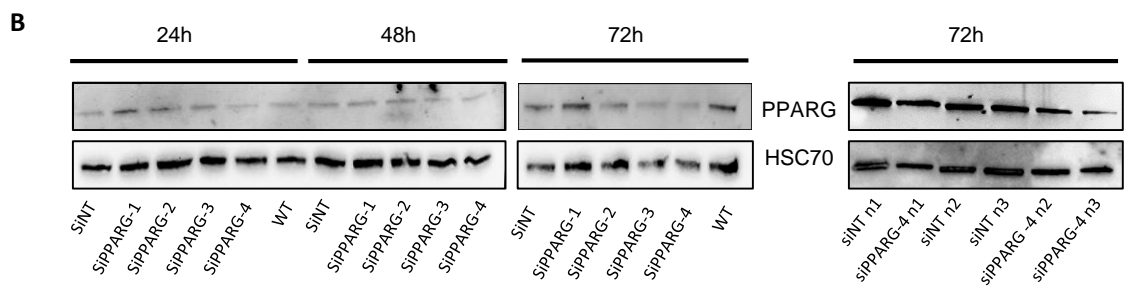

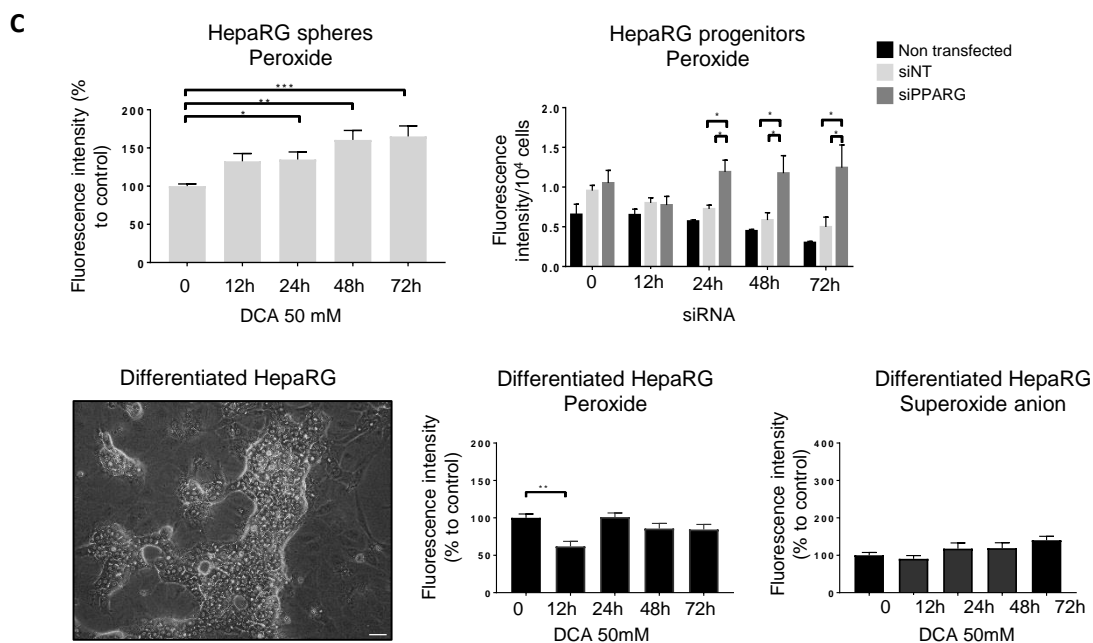

**Supplementary Figure 3**

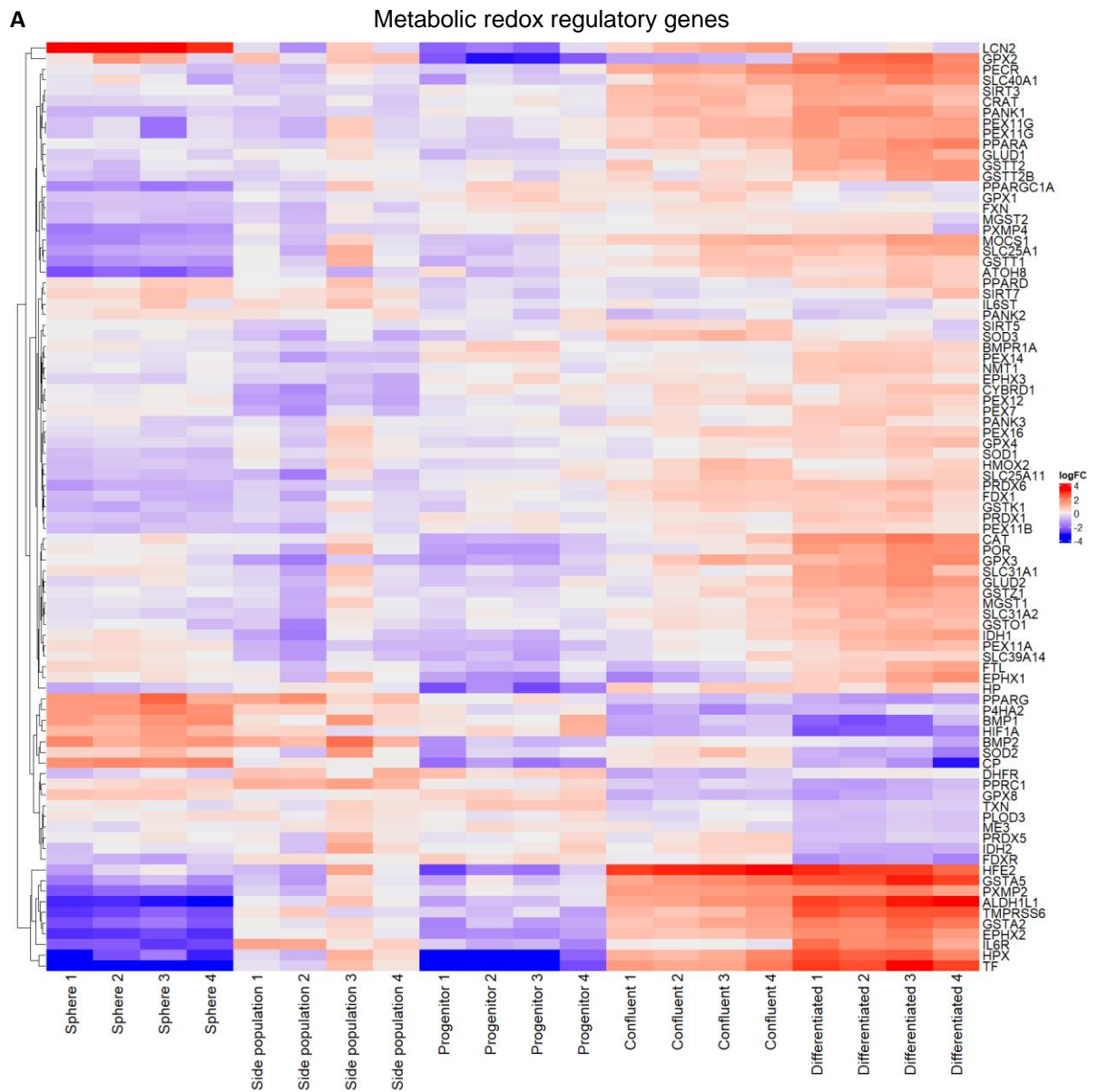

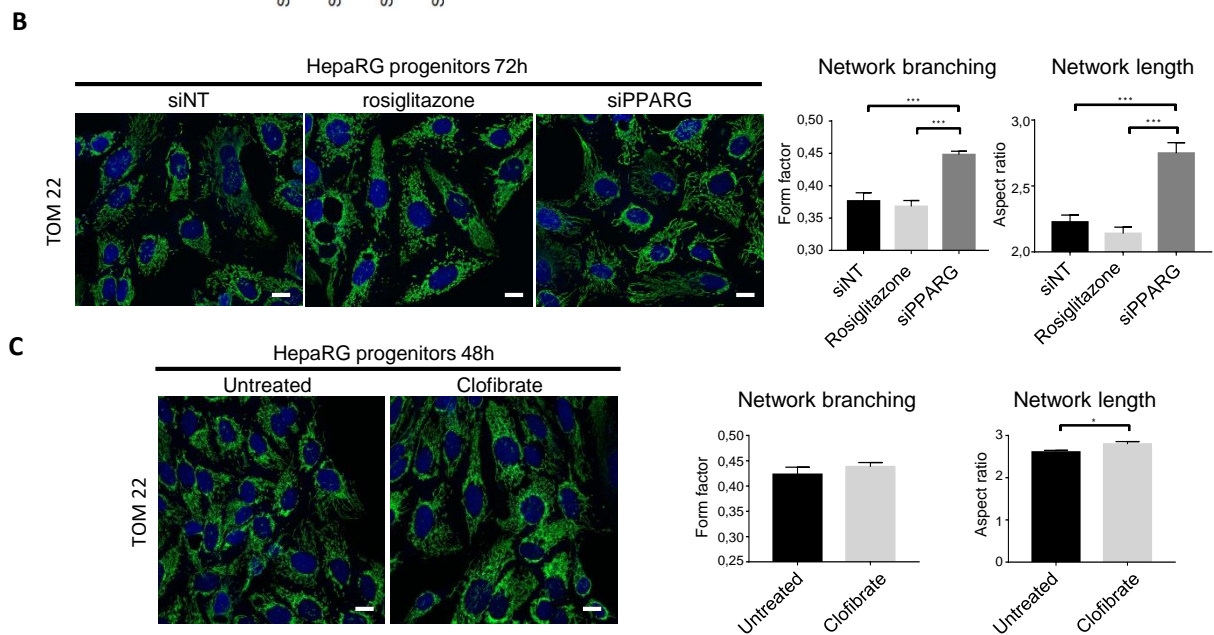

Supplementary Figure 4

#### Supplementary materials & methods

PPARγ, a key modulator of metabolic reprogramming, stemness and chemoresistance associated with retrodifferentiation in human hepatocellular carcinomas

Yoann Daniel<sup>1</sup>, Claudine Rauch<sup>1</sup>, Lucille Moutaux<sup>1</sup>, Karim Fekir<sup>1,2</sup>, Lise Desquilles<sup>1</sup>, Luis Cano<sup>1</sup>, Daniel Catheline<sup>1</sup>, Servane Pierre<sup>1</sup>, Agnès Burel<sup>3</sup>, Camille Savary<sup>1</sup>, Catherine Ribault<sup>1</sup>, Claude Bendavid<sup>1,4</sup>, Bruno Clément<sup>1</sup>, Caroline Aninat<sup>1</sup>, Vincent Rioux<sup>1</sup>, Orlando Musso<sup>1</sup>, Bernard Fromenty<sup>1</sup>, Florian Cabillic<sup>1,4#\*</sup> and Anne Corlu<sup>1,#\*</sup>

<sup>1</sup> Inserm, Univ Rennes, INRAE, NuMeCan Institute (Nutrition, metabolisms and cancer), Rennes, France

<sup>2</sup> Division of trauma and orthopaedic surgery, University of Cambridge, Addenbrooke's hospital, Hills road, Cambridge CB2 0QQ, UK

<sup>3</sup> Univ Rennes, CNRS, Inserm, Biosit UAR 3480 US\_S 018, MRIC-TEM platform, Rennes, France

<sup>4</sup> CHU of Rennes, Pontchaillou site, Rennes, France

### Florian Cabillic and Anne Corlu contributed equally to this work.

##### Data integration

Transcriptomes across several time points of mouse liver development (GSE90047(1) n = 21; GSE13149(2), n = 25) were quantile-normalized. Orthologs were obtained through the *gorth* function (*gProfiler* R package(3)). Batch effect was assessed and corrected with the ComBat algorithm(4) (*sva* R package (5)).

##### **Mitocarta gene set analysis**

The reads per million normalized gene expression data (n=374 HCCs) and clinical data were retrieved from the TCGA database. Genes from the Mitocarta\_up\_REST and Mitocarta\_dn\_REST signatures were considered as up-regulated or down-regulated, respectively. HCCs were classified as periportal, perivenous, ECM or STEM according to a hierarchical ascendant classification based on the Euclidean distance. PCA was computed using the *factoextra* and *FactoMineR* R packages and a biplot is plotted with confidence ellipses."

##### **Survival analysis**

Data from the Roessler (GSE14520, n = 247 HCCs) and ICGC (n = 232 HCCs) datasets were median-normalized using the DESeq2 R package. The reads per million normalized gene expression data (n=374 HCCs) from the TCGA database were also retrieved. The data relative to *PPARA*, *PPARG*, AFP levels and overall survival time and status were extracted and filtered for missing value. For each gene, patients were classified in "high expression" and "low expression" using the *surv\_cutpoint* function for the *survival* R package. Survival curves were fitted using the cutpoints of 2 genes at time as covariates producing 4 groups of patients. The differences in prognosis were evaluated according to Kaplan-Meier method. At the end, AFP levels were evaluated for each group to strengthen their prognostic values.

##### **Gene set enrichment analysis**

Gene set enrichment analysis (GSEA) was used to compare the 6115 genes differentially expressed (DEG) between immature and differentiating HepaRG cells with the Mitocarta 3.0 Gene Set (1136 genes).(6) Analysis was performed using the Java-tool developed at the Broad Institute (Cambridge, MA). 311 down-regulated genes in stem HepaRG were integrated into Enrichr website (https://maayanlab.cloud/Enrichr/).

##### **siRNA transfection**

The validated siRNA (LQ-003436-00-0002, On-TARGETplus Human PPARG 5468 siRNA – Set of 4, 2 nmol, Dharmacon) was used for knocking down the expression of *PPARG*. The ON-TARGETplus Non-targeting siRNA (D-001810-01-20, Dharmacon) was used as control. Transfection was performed using lipofectamine® RNAiMAX (Invitrogen, 13778-075). Briefly, one day after cell seeding, cells were incubated overnight with the liposome-DNA mix (1.5 µl RNAiMAX/25 pmoles DNA/105 cells) in Opti-MEM™ I Reduced Serum Medium (Gibco, 31985-070). Expression of *PPARG* was analyzed 48 and 72 hours after transfection by RT-qPCR. Knock down of *PPARG* in HepaRG sphere was performed by electroporation. Treatments with drugs were performed 48h after transfection for 48 hours. For N-Acetyl Cysteine (NAC, Merck, A9165) experiments, NAC was added to the medium at time of transfection and then every day.

##### **Drug treatments**

In absence of pre-treatment, adherent cells or spheres were cultured for 3 days in the presence of cisplatin (Mylan), sorafenib (Interchim, CJ592), dichloroacetate (Sigma-Aldrich, 347795-10G), clofibrate (Sigma Aldrich, C6643-5G), rosiglitazone (Sigma-Aldrich, R2408), SC-79 (4µg/ml, Tebu Bio, T2274) or LY294002 (10µM, Tebu Bio, B-0294) at the indicated concentration. For NAC and/or T0070907 (Tebu Bio, T6689) treatment, 24h after cell seeding, cells were pre-incubated for 24h with or without NAC (1mM) and/or T0070907 (10µM). Then, cells were cultured for 48h with or without NAC (1mM) and/or T0070907 (10µM) in the presence of cisplatin (Mylan), sorafenib (Interchim, CJ592) or dichloroacetate (Sigma-Aldrich, 347795-10G). NAC was added every day. To study the impact of co-treatment on HepaRG- and BC2-spheres, cells were pre-treated 24h with clofibrate at 100 or 500µM before 72h of co-treatment with cisplatin, sorafenib or dichloroacetate.

##### **Cytotoxicity and lipid peroxidation assays**

Cytotoxicity assays were performed using CellTiter-Glo® Luminescent Cell Viability Assay (Promega, G7571) on cells. Lipid peroxides that result in the formation of malondialdehyde (MDA) were measured as Thiobarbituric Acid Reactive Substances with the TBARS kit (Bio-Techne, KGE013) on HepaRG supernatants. Luminescence was measured by POLARstar Omega plate reader (BMG Labtech).

##### **Measurement of reactive oxygen species**

Cells were incubated with 5 μM Mitosox (ThermoFisher Scientific, M36008) or 2 μM H2DCFDA (Molecular Probes, D399) in warm Hank's balanced salt solution (HBSS) for 30 min at 37°C and 5% CO<sub>2</sub> in the dark. The medium was removed and the cells were washed with warm HBSS and the fluorescence intensity was measured by the POLARstar Omega plate reader (BMG Labtech). Two measurements were performed for each well with the two required wavelength pairs for Mitosox (ex520/em590) and DFCDA (ex485/em520). For probe control, specific wells containing cells were treated with 60 mM H<sub>2</sub>O<sub>2</sub> (Sigma-Aldrich, H1009) for 1 hour before probe incubation to validate probe activation.

##### **Real-time PCR**

Total RNA extraction and purification were performed using NucleoSpin RNA® Kit (Macherey-Nagel, 740955). Retrotranscription of RNA to cDNA was realized using High Capacity cDNA Reverse Transcription Kit (Applied Biosystems, 4368814). Total DNA extraction and purification were performed by DNeasy® Blood & Tissue Kit (Qiagen, 69504). Real-time qPCR was performed with Thermocycler StepOnePlus™ device (Applied Biosystems) using SYBR Green PCR Master Mix (Applied Biosystems, 4309155). Gene expression was normalized by housekeeping gene Tata Binding Protein for (*TBP*) cDNA and Ribosomal Protein S6 (*RPS6*) for DNA. The primer sequences are listed in the following table.

| Primer | Gene name | Forward | Reverse |
| --- | --- | --- | --- |
| TBP | TATA-Box Binding Protein | 5'-ACT-CCA-CTG-TAT-CCC-TCC-CC-3' | 5'-CAG-CAA-ACC-GCT-TGG-GAT-TA-3' |
| ALB | Albumin | 5'-TGCTTGAATGTGCTGATGACAGG-3' | 5'-AAGGCAAGTCAGCAGGCATCTCATC-3' |
| CD44 | CD44 Molecule | 5'-GGC-TTT-CAA-TAG-CAC-CTT-GC-3' | 5'-CAC-GTG-CCC-TTC-TAT-GAA-CC-3' |
| FAT/CD36 | Fatty Acid Translocase | 5'-GCC-AGG-TAT-TGC-AGT-TCT-TTT-C-3' | 5'-TGT-CTG-GGT-TTT-CAA-CTG-GAG-3' |
| PPARG | Peroxisome Proliferator Activated Receptor Gamma | 5'-AAG-GCC-ATT-TTC-TCA-AAC-GA-3' | 5'-AGG-AGT-GGG-AGT-GGT-CTT-CC-3' |
| PPARA | Peroxisome Proliferator Activated Receptor Alpha | 5'-GTT-CTG-GAA-GCT-TTG-GCT-TTA-C-3' | 5'-GAA-AGC-GTG-TCC-GTG-ATG-A-3' |
| PPARD | Peroxisome Proliferator Activated Receptor Delta | 5'-AGC-ATC-CTC-ACC-GGC-AAA-G-3' | 5'-CCA-CAA-TGT-CTC-GAT-GGC-AGG-CG-3' |
| PLIN2 | Perilipin 2 | 5'-GCT-CCA-TTC-TAG-TGT-TCA-CCT-G-3' | 5'-CTC-CTT-TTC-CAC-TCT-ACC-CAT-G-3' |
| PDK4 | Pyruvate Dehydrogenase Kinase 4 | 5'-CTCGCGCTAGAGCCCG-3' | 5'-GCATTTTCTGAACCAAAGTCCAG-3' |
| ANGPTL4 | Angiopoietin Like 4 | 5'-GACCAAGGGGCATGGAGCTT-3' | 5'-CAGGGGACCTACACACAACAG-3' |
| KLF4 | Kruppel Like Factor 4 | 5'-GAC-GCT-GCT-GAG-TGG-AAG-AG-3' | 5'-AGA-CAA-TCA-GCA-AGG-CGA-GT-3' |
| CDH1 | Cadherin 1 | 5'-AGT-GGG-CAG-AGA-TGG-TGT-GA-3' | 5'-TAG-GTG-GAG-TCC-CAG-GCG-TA-3' |
| RPS6 | Ribosomal Protein S6 | 5'-TGA-TGT-CCG-CCA-GTA-TGT-TG-3' | 5'-TCT-TGG-TAC-GCT-GCT-TCT-TC-3' |

#### Protein extraction and western blot analysis

Cells were lysed in ice-cold RIPA buffer containing phosphatase and protease inhibitors (ThermoFisher Scientific, A32959). Protein concentration was determined using the Pierce BCA Protein Assay Kit (Thermo Fisher Scientific, 23227) and equal amounts of protein were diluted with NuPAGE LDS Sample Buffer (Thermo Fisher Scientific, NP0007). Samples were then separated by SDS-PAGE using iD PAGE Gel 4–12% (Eurogentec, ID-PA4121-012). Proteins were transferred with iBlot Gel Transfer Stacks Nitrocellulose (Life Technologies, IB301001). Membranes were blocked in 3% BSA in TBST (10 mM Tris-HCl, 100 mM NaCl, 0.02% Tween 20) for 1h at room temperature and incubated overnight at 4 C with antibodies against PPARγ (Santa Cruz Biotechnology Cat# sc-7196, RRID:AB\_654710), PPARα

(Invitrogen cat# MA5-37652, RRID:AB\_2897578 ) or HSC70 (Santa Cruz Biotechnology Cat# sc-7298, RRID:AB\_627761). Membranes were washed with TBST, incubated for 1h at room temperature with appropriate HRP secondary antibodies (Agilent Cat# P0447, RRID:AB\_2617137, and Cat# P0448, RRID:AB\_2617138, respective dilutions 1:5000 and 1:10,000), washed with TBST, and then visualized by enhanced chemiluminescence (Pierce ECL Western Blotting Substrate; ThermoFisher Scientific) using Fusion FX imaging system (Vilber Lourmat). Protein content was quantified by densitometry with ImageJ software (National Institutes of Health, Bethesda, MD).

##### **Mitochondrial mass by flow cytometry**

Cells were incubated with 100 nM of MitoTracker® Green FM (ThermoFisher Scientific, M7514) for 30 min in HBSS (Gibco, 14175095). Cells were then washed in HBSS, detached with 0.05% trypsin (Gibco, 25300-054), resuspended in HBSS and ran through the LSR Fortessa X-20 flow cytometer.

##### **Dosage of pyruvic and lactic acids**

Supernatants of culture were collected and put into a tube containing 2 ml of perchloric acid. The medium had been renewed 24 hours before this sampling. Pyruvic acid and lactic acid determinations were performed by photometry using a COBAS c111 (ROCHE) in the biochemistry/toxicology laboratory of the University Hospital of Rennes.

##### **Assessment of neutral lipids with Nile Red**

Cells were washed with PBS, fixed and stained with PBS containing 4% formaldehyde and 10 µg/mL Hoechst 33342 dye for 30 min and washed 3 times with PBS. Cells were then incubated with PBS containing 0.1 µg/ml Nile Red (ThermoFisher Scientific, N1142) for 30 min and washed once. Fluorescence intensity was measured by the POLARstar Omega plate reader (BMG Labtech). Neutral lipids were then normalized per number of nuclei and expressed relative to control cells.

##### **Lipid extraction, lipid species separation and fatty acid analysis**

Total lipids were extracted twice from cells with hexane/isopropanol (3/2 v/v), after acidification with HCl 3M, as previously described.<sup>(7)</sup> Lipid species were then separated by thin-layer chromatography (TLC) using silica gel H plates (0.5 mm thickness) and a mixture of hexane:diethylether:acetic acid (85:15:1 v/v/v) for development, after addition of internal standards (diheptadecanoylphosphatidycholine, heptadecanoic acid, triheptadecanoylglycerol and cholesteryl heptadecanoate).<sup>(8)</sup> Phospholipids (PL), free fatty acids (FFA), triglycerides (TG) and cholesterol esters (CE) were scraped off the plates and extracted with 2 mL of methanol. Total lipids and lipid species were converted to Fatty Acid Methyl Esters (FAMES), by successive saponification (1 mL of 0.5 M NaOH in methanol at 70°C for 30 min) and methylation (1 mL of BF<sub>3</sub> 14% in methanol at 70°C for 30 min).<sup>(9)</sup> Gas chromatography-mass spectrometry (GC-MS) analysis of FAMES was subsequently performed using an Agilent Technologies 7890A GC system (Agilent, Les Ulis, France) with a bonded silica capillary column (BPX 70, 60 m × 0.25 mm; SGE, Melbourne, Australia) containing a polar stationary phase of 70% cyanopropyl polysilphenylene-siloxane (0.25 µm film thickness). Helium was used as carrier gas (average velocity 24 cm/s). The column temperature program started at 150°C and gradually increased at 4°C/min to 250°C, and held at 250°C for 10 min. Mass spectra were recorded with an Agilent Technologies 5975C inert MSD with triple axis detector.<sup>(10)</sup> The mass spectrometer was operated under electron impact ionization conditions (electron energy 70 eV, source temperature 230°C). Data were obtained in the full scan mode with a mass range of m/z 50–550 atomic mass units (amu). Peak integration was performed with

MassHunter Workstation Software Qualitative Analysis Version B.07.00 for Windows (Agilent Technologies). Identification of the FAMES was based upon retention times (Rt) obtained for methyl ester of authentic standards, when available. The National Institute of Standards and Technology database (NIST version 2.2) was used to identify unknown fatty acids. All identified fatty acids with a signal/noise above 10 were considered in the analysis. Results were expressed as the mass of identified fatty acids/cell number.

##### **Electron microscopy**

Cells were fixed with 2.5% glutaraldehyde (Sigma-Aldrich, G7526) and treated by 1% osmium (Electron Microscope Sciences, 19150) in 0.2M sodium cacodylate buffer pH=7.2 (Electron Microscope Sciences, 11650). The dehydration was carried out by successive alcohol baths (increasing alcohol content: 50%, 70%, 80%, 90%, 100%) followed by twice impregnation with an Epon / DMP30 mixture (Sigma-Aldrich, 45348) and placed at 37°C. The next day, capsules were inserted into the same resin and the plates were placed at 37°C. Two days later, the capsules were filled with an Epon/DMP30 mixture and placed in an oven at 60°C to polymerize for 24 hours. The capsules were detached from the plates and cut with an ultra-microtome (UC7, Leica). Sections were stained with uranyl acetate and the images acquired with a Gatan Orius SC1000 wide angle camera on a JEOL JEM-1400 electron microscope. For each cell type, the TEM experiments were carried out at least twice. At least two capsules were cut by experiment and for each section examined by TEM, 4 different fields were investigated in order to overview the cells layer.

##### **Confocal microscopy**

HepaRG cell line was cultured in compartmented Lab-Tek™. After culture, cells were fixed with 4% paraformaldehyde (ThermoFisher Scientific, 28908) and permeabilized by PBS with

1% BSA (Sigma-Aldrich, A2153) and 0.5% saponin (Sigma-Aldrich, 47036) for 1h at room temperature. First incubation was done with antibody against mitochondrial import receptor subunit (TOM22) at 1/5000 (Abcam Cat# ab57523, RRID:AB\_945897) in PBS with 1% BSA and 0.5% saponin overnight at 4°C. Next day, compartmented slats were washed with PBS and secondary anti-mouse antibody coupled with green FITC (Jackson ImmunoResearch Labs Cat# 115-095-003, RRID:AB\_2338589) for 1 hour at room temperature, followed by PBS washing and Hoescht 33342 labelling for 15 min. The cells were observed with a LEICA DMI 6000 CS confocal microscope. The acquisition of images was performed with the LAS AF software. Image analysis was performed with ImageJ software, based on the work of Koopman et al.(11)

##### **Immunohistochemistry**

Tissue Microarray (TMA) comprises 25 HCC patients and 5 histologically normal livers, all in triplicate as previously described.(12) Briefly, TMA construction was done with a MiniCore3 tissue arrayer (Alphelys, France). Five-µm microtome sections were processed for immunohistochemistry with a Discovery XT from Ventana Medical Systems (Roche) slide staining system. PPARG antibodies used were purchased from Invitrogen (Cat# MA5-14889, RRID:AB\_10985650). Stained slides were converted into high-resolution digital data with a NanoZoomer digital slide scanner (Hamamatsu, France). Digital slides were viewed using NDP.view software (Hamamatsu).
